# Presynaptic dysfunction in *CASK*-related neurodevelopmental disorders

**DOI:** 10.1101/863308

**Authors:** Martin Becker, Francesca Mastropasqua, Jan Philipp Reising, Simon Maier, Mai-Lan Ho, Ielyzaveta Rabkina, Danyang Li, Janina Neufeld, Lea Ballenberger, Lynnea Myers, Viveka Moritz, Malin Kele, Josephine Wincent, Charlotte Willfors, Rouslan Sitnikov, Eric Herlenius, Britt-Marie Anderlid, Anna Falk, Sven Bölte, Kristiina Tammimies

## Abstract

*CASK*-related disorders are a genetically defined group of neurodevelopmental syndromes. There is limited information about the effects of *CASK* mutations in human neurons. Therefore, we sought to delineate *CASK* mutation consequences and neuronal level effects using induced pluripotent stem cell-derived neurons from two mutation carriers; one male diagnosed with ASD and a female with MICPCH. We show a reduction of the CASK protein in maturing neurons from the mutation carriers, which leads to significant downregulation of gene sets involved in presynaptic development and CASK protein interactors. Furthermore, *CASK*-deficient neurons showed decreased inhibitory presynapse size as indicated by VGAT staining, which may alter the excitatory-inhibitory (E/I) balance in developing neural circuitries. Using *in vivo* magnetic resonance spectroscopy quantification of GABA in the male mutation carrier, we further highlight the possibility to validate *in vitro* cellular data in brain. Our data shows that future pharmacological and clinical studies on targeting presynapses and E/I imbalance could lead to specific treatments for *CASK*-related disorders.

**Highlights:** Modelling of CASK-related disorders using iPSC-derived human neuronal cells

*CASK* mutations cause dysregulation of its protein interactor partners

Reduced CASK levels primarily affect inhibitory presynapse development

*In vitro* GABAergic phenotype predicts *in vivo* neurotransmitter levels

## Introduction

The identification of genetic variants underlying neurodevelopmental disorders (NDDs), such as intellectual disability (ID), autism spectrum disorder (ASD) and attention deficit hyperactivity disorder (ADHD) as well as epilepsies has increased at a rapid pace in recent years with high level of pleiotropy across the conditions (De Rubeis et al., 2014; Deciphering Developmental Disorders, 2017; Plummer et al., 2016; Satterstrom et al., 2019). Findings indicate shared molecular mechanisms underlying the diverse clinical phenotypes. However, for many of the identified risk variants and genes, the molecular and neuronal outcomes are not well understood. One pleiotropic gene is the *calcium/calmodulin-dependent serine protein kinase* (*CASK*), located on chromosome Xp11.4, in which pathogenic variants underlie a range of NDDs. Genetic variants in *CASK* were first described in cases with microcephaly with pontine and cerebellar hypoplasia (MICPCH), followed by the identification in cases with X-linked ID (XL-ID), developmental delay (DD) and ASD (Deciphering Developmental Disorders, 2017; Iossifov et al., 2014; Moog et al., 2015; Moog et al., 2011). The majority of cases reported with *CASK*-related disorders are females with MICPCH, caused by heterozygous loss-of-function (LoF) variants (Burglen et al., 2012; Hayashi et al., 2017; Moog et al., 2011; Najm et al., 2008). Skewed X-chromosome inactivation (XCI) has shown protective effects against more severe phenotypes (Moog et al., 2011; Seto et al., 2017). Missense variants, found both in females and males cause microcephaly, XL-ID, DD or ASD (Cristofoli et al., 2018; Deciphering Developmental Disorders, 2017; Gupta et al., 2014; Hackett et al., 2010; Iossifov et al., 2014; LaConte et al., 2018; Moog et al., 2015; Piluso et al., 2009; Sanders et al., 2012; Seto et al., 2017). The molecular pathways and cellular phenotypes associated with genetic variants causing *CASK*-related disorders are mostly unknown, especially in human neurons.

CASK is ubiquitously expressed with high expression in the developing human brain (Stevenson et al., 2000). The protein structure consists of scaffolding domains L27, PDZ, and SH3, as well as a Ca^2+^/calmodulin-dependent protein kinase and guanylate kinase domain (Hata et al., 1996). In neurons, CASK is involved in pre- and postsynaptic signaling. At the presynapse, CASK regulates the synaptic vesicle exocytosis and neuronal cell adhesion through a tripartite complex with VELI1 (*LIN7A)* and MINT1 (*APBA1)* and direct interaction with NRXN1 (*NRXN1)* (Butz et al., 1998; LaConte et al., 2016; Sudhof, 2012). The tripartite protein complex VELI1-MINT1-CASK is unaffected by XL-ID and MICPCH missense mutations (LaConte et al., 2014; LaConte et al., 2018). Instead, the CASK-NRXN1 interaction is disrupted, suggesting that the absence of this interaction is associated with the MICPCH phenotype. In the postsynaptic density, CASK contributes to the regulation of ionotropic receptor trafficking (Lin et al., 2013). Electrophysiological assessment of CASK negative neurons from mice showed a shift in excitatory-inhibitory (E/I) balance with increased spontaneous miniature excitatory postsynaptic currents (mEPSC) and decreased miniature inhibitory postsynaptic currents (mIPSC) (Atasoy et al., 2007; Mori et al., 2019). Interestingly, *Nrxn1* KO mouse neurons and human neuronal models with *NRXN1* LoF mutations have elevated *CASK* levels and decreased mEPSCs (Pak et al., 2015). In addition to the role at the synapse, CASK has been shown to act as a co-regulator of transcription through TBR1 (*TBR1*), CINAP (*TSPYL2*) and BCL11A (*BCL11A*) (Hata et al., 1996; Kuo et al., 2010; Wang et al., 2004), all of which have been implicated in NDDs (Deriziotis et al., 2014; Dias et al., 2016; Moey et al., 2016).

Here, we elucidate the consequences of *CASK* pathogenic variants using human induced pluripotent stem cell-derived neuronal models from two mutation carriers, one diagnosed with ASD and one with MICPCH. Our transcriptional, morphological, and functional analyses reveal that reduced levels of CASK are sufficient to induce aberrant presynaptic development, and have wide effects on its interaction network, as well as on multiple developmental pathways. In addition, for the first time we explore the possibility to validate neuronal findings from iPSC models using *in vivo* brain data. In conclusion, our data suggest that *CASK*-related disorders are synaptopathies that affect the E/I balance in maturing neural circuits.

## Results

### CASK variants lead to mutant transcripts and reduced CASK_WT_ expression

To analyze the converging molecular and cellular consequences of pathogenic variants affecting *CASK* across the associated NDD spectrum, we recruited two families with unique *CASK* genetic alterations, representing ASD and MICPCH phenotypes. First, we identified a male individual diagnosed with ASD carrying a *CASK* mutation from the Roots of Autism and ADHD Twin Study in Sweden (RATSS) (Bölte et al., 2014). He had a primary diagnosis of ASD and general intellectual abilities in the upper average range without cognitive impairments (total IQ=121) and additional diagnoses of ADHD and tic disorder (Figure 1A, Table S1). His monozygotic co-twin was diagnosed with ASD prior to the RATSS study. However, the co-twin did not fulfill clinical consensus ASD diagnostic criteria at re-assessment during the study, but still exhibited autistic symptoms (Table S1). Anatomical magnetic resonance imaging (MRI) scans of both twins revealed mild cerebral atrophy and mild cerebellar hypoplasia within normal variation (Figure S1A, B). No clinically significant copy number variants were found in the twin pair. Rare variants from the exome sequencing were prioritized based on variant effect, inheritance mode, and genes involved in NDDs. We identified a novel maternally inherited splice-site variant in intron 14 of the *CASK* gene in both twins (chrX: 41,586,906 C>A (hg38); NM_001126055: c.1296+1 G>T, Figure S1C). The variant disrupts the consensus splice donor site at +1 G nucleotide (Figure S1D). *CASK* is intolerant to LoF mutations (pLI=1), and three putative LoF splice site mutations reported in the general population are outside the main transcript or tissue-specific transcripts, as reported by the GTEx consortium (Carithers et al., 2015; Karczewski et al., 2019). Additionally, both twins carry a *de novo* heterozygous missense mutation affecting *DIAPH1* exon 15 (chr5: 141,574,906 C>A (hg38); NM_005219: c.1528 G>A: p.E510K, Figure S1E). Heterozygous truncating mutations of this gene are reported in cases of congenital deafness (Lynch et al., 1997; Neuhaus et al., 2017; Stritt et al., 2016), and homozygous LoF mutations are linked to microcephaly (Al-Maawali et al., 2016; Ercan-Sencicek et al., 2015). None of the reported dominant mutations in *DIAPH1* have been linked to NDDs. Of interest, slight hearing problems in one ear were reported for one of the twins (Table S1).

**Figure 1.**
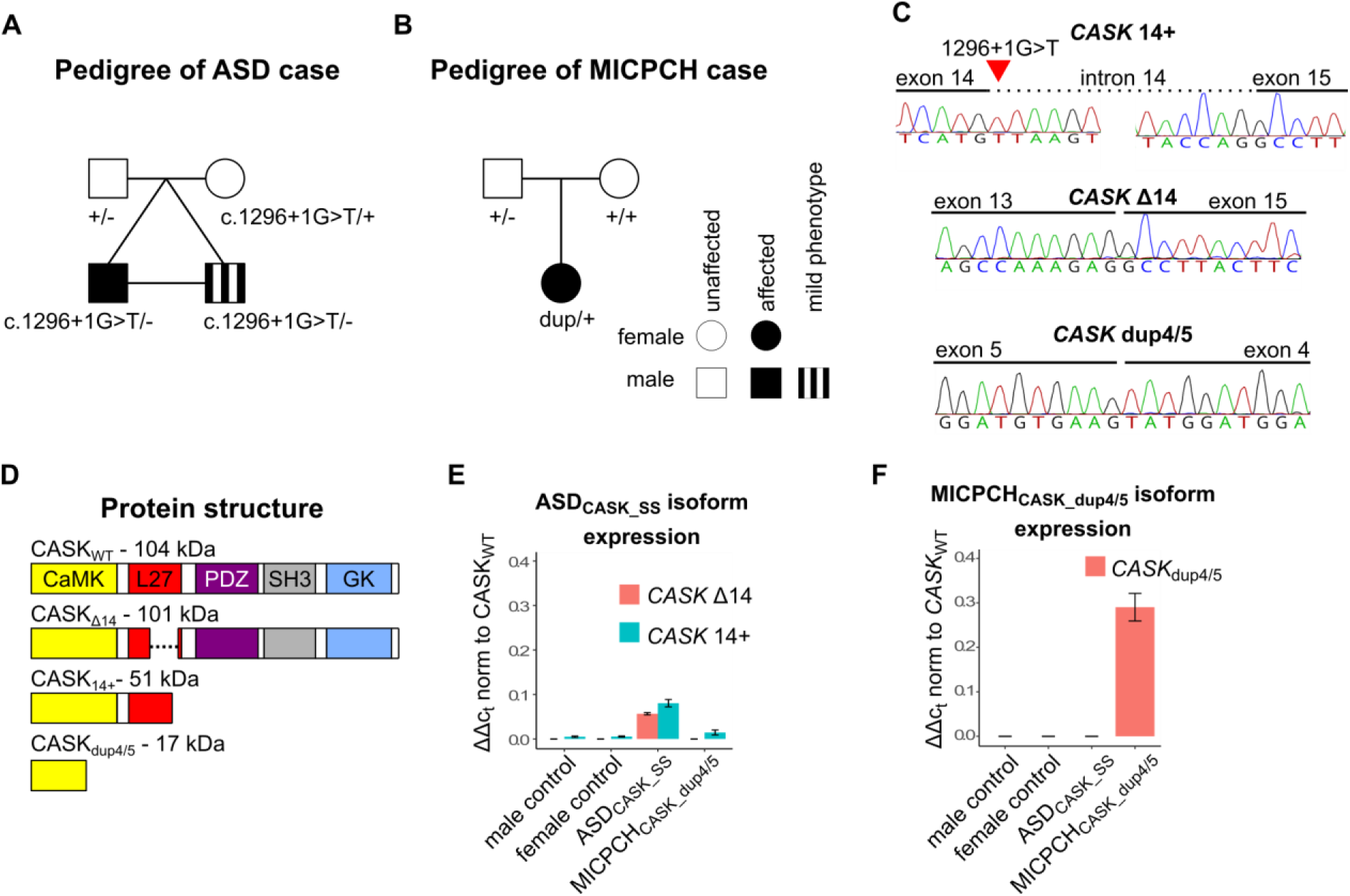
*CASK* mutations affect wild type expression in carriers. (A) Pedigree of a male diagnosed with ASD and his monozygotic co-twin who exhibit autistic traits, both carrying a splice site mutation in the X-linked *CASK* gene (chrX: 41,586,906 C>A (hg38); NM_001126055: c.1296+1 G>T) inherited from a typically developed mother. (B) Pedigree of a female diagnosed with MICPCH, carrying a *de novo* duplication of 54.9kb surrounding exons 4 and 5 (chrX:4217380-41765176 (hg38)) of the *CASK* gene. (C) Sanger sequence of *CASK* mRNA isoforms in cDNA of ASD_CASK_SS_ fibroblasts and MICPCH_CASK_dup4/5_ NES cells. (D) Protein domain structure of CASK_WT_ and *in silico* predicted domain structure of CASK_Δ14_, CASK_14+_ and CASK_dup4/5_. (E) RT-qPCR quantification of *CASK*_Δ14_ and *CASK*_14+_ expression in three biological replicates of case and control NES cells. (F) RT-qPCR quantification of *CASK*_dup4/5_ in three biological replicates of case and control NES cells.

We also recruited a female with a classical presentation of MICPCH syndrome, caused by a *de novo* duplication of 54.9 kb at chromosome Xp11.4, spanning two exons of *CASK* (Wincent et al., 2015) (Figure 1B). Her degree of ID is in accordance with earlier described individuals with MICPCH. She did not communicate verbally, had pontocerebellar hypoplasia, scoliosis, optic nerve hypoplasia, and infantile spasms and epilepsy. Wincent *et al*. provided detailed description of the individual (referred to as patient seven in their study), including clinical symptoms, structural brain MRI and genetic analysis.

To investigate the consequences of the identified splice-site and duplication variants to *CASK* mRNA, we obtained fibroblasts from the two mutation carriers hereafter referred to as ASD_CASK_SS_ and MICPCH_CASK_dup4/5_. The obtained fibroblasts were transformed into induced pluripotent stem cells (iPSCs). The iPSCs had pluripotent marker expression, normal karyotype, and matched donor genotypes (Figure S1F, S1G). Furthermore, the iPSCs were differentiated to long-term self-renewing neuroepithelial-like stem cells (further referred to as NES cells, Figure S1H) (Falk et al., 2012). NES cell lines derived from two sex-matched typically developed individuals were obtained for controls (Falk et al., 2012; Uhlin et al., 2017). We detected two mutant *CASK* mRNA isoforms in the NES cells from ASD_CASK_SS_ (Figure 1C). One isoform lacks exon 14 (CASK_Δ14_), and one isoform retains 196 nucleotides of intron 14 (CASK_14+_), resulting in a premature stop codon (Figure 1D). Although ASD_CASK_SS_ is hemizygous, we detected wild-type *CASK* mRNA (CASK_WT_) with intact joining of exon 14 and 15 constituting majority of the expression. A minor proportion of *CASK* mRNA consisted of mutant CASK_Δ14_ and CASK_14+_ isoforms with approximately 6% and 8%, respectively (Figure 1E). CASK_14+_ isoform was detected at low levels in control and MICPCH samples, presenting likely pre-mRNA. The presence of CASK_WT_ in the cells of ASD_CASK_SS_ indicate rescue mechanisms for normal splicing possibly through mRNA editing. The MICPCH_CASK_dup4/5_ NES cells expressed an mRNA isoform with a tandem duplication of exons 4 and 5 (Figure 1C), which leads to a frame shift and early stop codon in the *CASK* coding sequence (Figure 1D). This mutant isoform constitutes ∼29% of total *CASK* mRNA in MICPCH_CASK_dup4/5_ cells (Figure 1F). In summary, we show that NES cells from both mutation carriers express CASK_WT_ and detectable levels of mutant mRNA isoforms that may interfere with CASK function if translated to protein.

### Reduction of CASK expression in maturing neurons from mutation carriers

Next, we differentiated NES cells to neurons and investigated the neurodevelopmental expression of *CASK* variants *in vitro* (Figure S1H). To cover stages of early neuronal differentiation, we collected RNA and protein from 8, 16, and 28 days of neuronal differentiation. None of the predicted mutant protein isoforms were detected from capillary western blotting using the N-terminal antibody (Figure S2A), suggesting that mutant mRNAs are removed by nonsense-mediated decay (NMD). *CASK* mRNA and protein expression increased with differentiation in all cell lines (Figures 2A, B). However, at 28 days of differentiation, the ASD_CASK_SS_ cells expressed significantly reduced levels of CASK mRNA and protein in comparison to both controls (p=0.02 and p=0.01, respectively, ANOVA posthoc Tukey). mRNA but not protein levels were increased in the MICPCH_CASK_dup4/5_ from NES stage to day 16 and leveled with controls on day 28 (day 0 p<0.001, day 8 p<0.001, day 16 p=0.001, day28 p=1, ANOVA posthoc Tukey). The protein levels at day 28 are reduced in comparison to the sex-matched female control (p=0.04, ANOVA posthoc Tukey, Figure S2B). In addition to NMD, biallelic expression from both X-chromosomes may explain the discrepancy between increased CASK mRNA and normal protein levels in MICPCH_CASK_dup4/5_.

**Figure 2.**
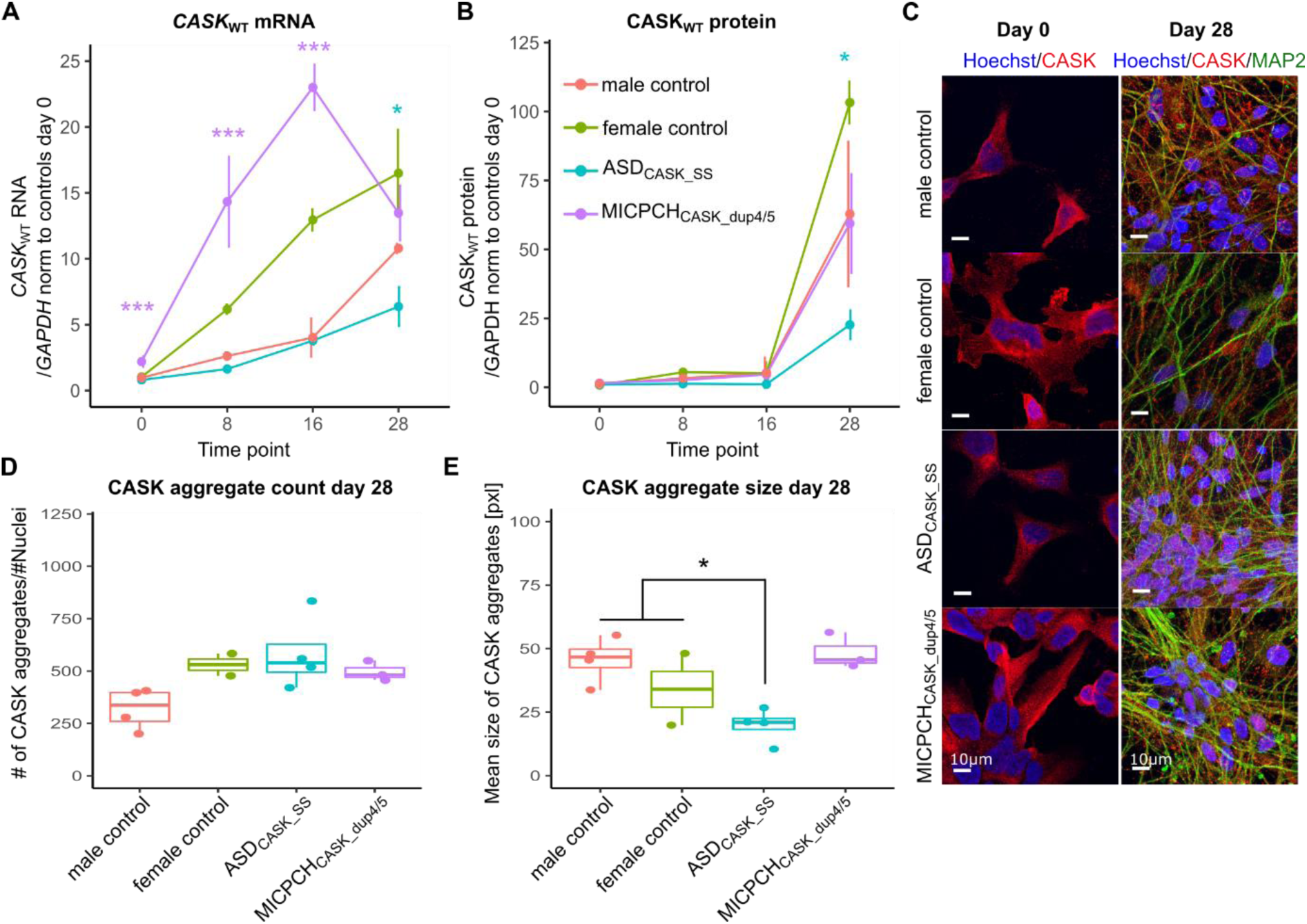
Reduced protein levels decrease CASK aggregate size in differentiating neurons. (A) RT-qPCR quantification of *CASK*_WT_ in three biological replicates of differentiating neurons. (B) Protein quantification of CASKWT protein in two biological replicates of differentiating neurons using capillary western blot quantification. (C) Representative confocal microscopy images of NES cells (day 0) and differentiated neurons (day28) immunostained for CASK (red), MAP2 (green) and Hoechst (blue). Scale = 10µm. (D and E) Quantification of CASK aggregate (D) number and (E) size from confocal images. Female control (n=2), male control (n=4), ASDCASK_SS (n=4) and MICPCHCASK_dup4/5 (n=3). Statistical differences between cases and controls were calculated using ANOVA with post-hoc Tukey HSD. *p < 0.05, ***p < 0.001. Asterisks in A and B are colour-coded according to case cell lines.

We further investigated the cellular localization of CASK protein in the ASD_CASK_SS_ and MICPCH_CASK_dup4/5_ cells using the N-terminal binding antibody. In NES cells, we observed small CASK aggregates in the cytoplasm of all cell lines with no aberrant nuclear staining (Figure 2C, day 0). With continuing differentiation, the CASK aggregates remained in the cytoplasm, including the neurites (Figure 2C, day 28) with comparable localization in all cell lines. In MICPCH_CASK_dup4/5_, CASK staining was detected in all cells demonstrating that the *CASK*_WT_ allele is active in all cells. Quantification of CASK aggregate size and number showed that aggregates were significantly smaller in differentiated ASD_CASK_SS_ neurons compared to both controls (p=0.019, ANOVA posthoc Tukey, Figure 2D), with similar aggregate numbers per nuclei (p=0.12, ANOVA posthoc Tukey, Figure 2E). However, we did not observe significant differences in MICPCH_CASK_dup4/5_ neurons compared to controls.

### X-chromosome activity in MICPCH_CASK_dup4/5_ cells

Random XCI were tested in blood from the MICPCH_CASK_dup4/5_ case in clinical setting showing a relation of 24% and 76%, consistent with random inactivation. We hypothesized that the discrepancy between CASK mRNA and protein expression during differentiation was due to biallelic expression from wild type and mutant alleles. To investigate XCI and escape of *CASK* mRNA in our model, we performed SMART-Seq2 single-cell RNA-sequencing on MICPCH_CASK_dup4/5_ neurons differentiated for 28 days. We obtained quality reads for 383 cells and identified the CASK_dup4/5_ specific exon5-exon4 junction in 22 cells (Figure 3A, Figure S3C). We compared this detection rate to the detection of the adjacent exons 4 (62 cells) and 5 (51 cells), indicating that 35% to 43% of cells expressed CASK_dup4/5_. The frequency of CASK_dup4/5_ expressing cells is indicative of random XCI and comparable to detection levels of the mutant isoform in RT-qPCR (Figure 1F). The presence of wild type protein in all MICPCH_CASK_dup4/5_ cells hint towards the escape of *CASK*_WT_ from XCI.

**Figure 3.**
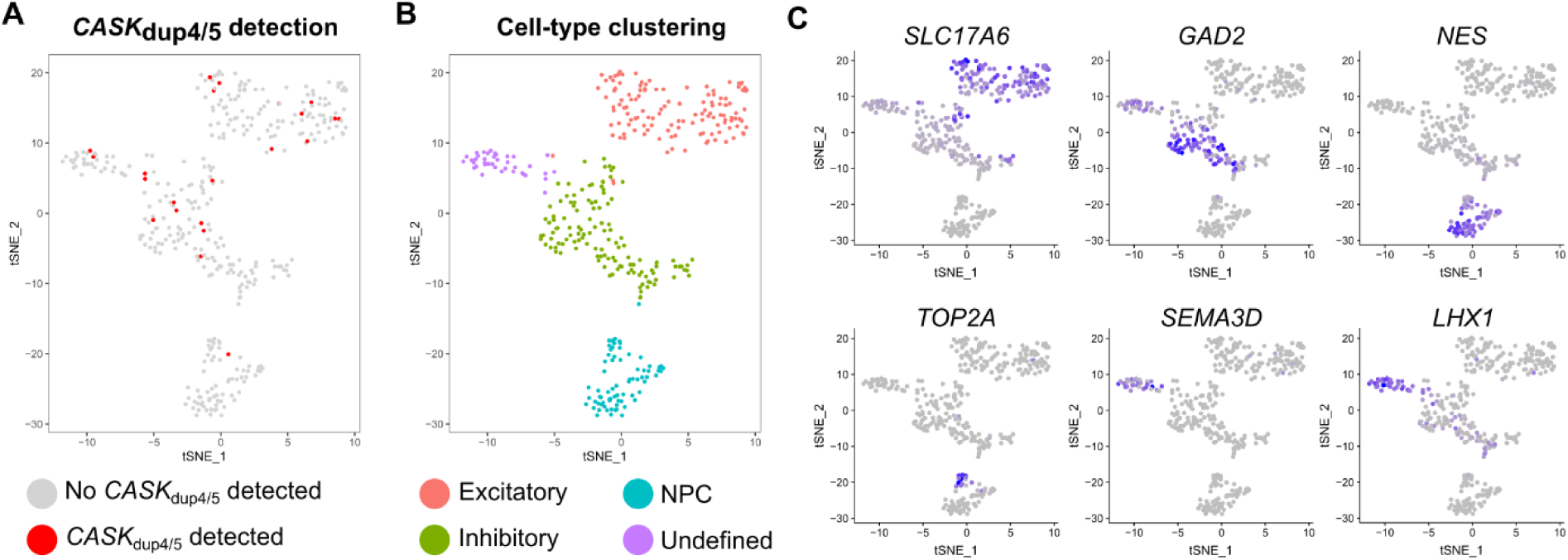
Single cell transcriptomics of MICPCH_CASK_dup4/5_. (A-C) tSNE scatterplot of 383 MICPCH_CASK_dup4/5_ cell transcriptomes after 28 days of differentiation coloured by (A) the detection of mutant *CASK*_dup4/5_ expression, (B) the identified cell type clusters and (D) expression of *SLC17A6*, *GAD2*, *NES*, *TOP2A*, *SEMA3D* and *LHX1*.

Moreover, we identified cell types in the differentiated neuronal culture from the single-cell transcriptomes and mapped *CASK*_dup4/5_ expression. The cultured cells clustered into four neuronal subtypes, which we identified as neural progenitor cells (NPC), excitatory and inhibitory neurons, as well as one cluster of undefined maturing neurons (Figure 3B). Excitatory neurons express the *SLC17A6* gene encoding for the vesicular glutamate transporter VGLUT2, and inhibitory neurons express the *GAD2* gene encoding the GAD65 enzyme required for GABA production (Figure 3C). NPCs express neuronal progenitor marker Nestin (*NES)* and cell-cycle gene *TOP2A*. Selective markers of the fourth cluster included genes involved in axon guidance *SEMA3D* and *LHX1*. As expected from the increase of CASK expression during differentiation, we observed higher expression in differentiating neurons compared to NPCs (adj.p=6E-6, DESeq2, Figure S3D). Expression of the mutant RNA isoform does not cluster within one cell type (Figure 3A) indicating that CASK protein expression changes are not cell type specific.

### Reduced CASK levels alter presynaptic development

We performed bulk RNA-sequencing of day 28 maturing neurons from all cell lines to identify *CASK*-dependent expression changes. The RNA-sequencing data showed a similar distribution of *CASK* expression for ASD_CASK_SS_ and MICPCH_CASKdup4/5_ as earlier detected by RT-qPCR (Figure S4A). We also quantified the isoform-specific reads from exon-exon spanning fragments (Table S2). Similar to RT-qPCR results at NES stage, we detected a low abundance of *CASK* mutant isoforms *CASK*_14+_ and *CASK*_Δ14_ of 4% and 8% in ASD_CASK_SS_, respectively. In addition, we detected both the mutant and reference nucleotide in *CASK*_14+_ reads that span the mutation site (Figure S4B). The *de novo* variant in *DIAPH1* was present in around 50% of *DIAPH1* transcripts (Figure S4C). In the MICPCH_CASK_dup4/5_ transcriptome, we detected 14% *CASK*_dup4/5_ specific sequence reads aligning to the exon 5-exon 4 junction. MICPCH_CASK_dup4/5_ cells also showed downregulation of the XCI regulating transcripts *XIST* and *TSIX*, which could explain activation of biallelic *CASK* expression (Figure S4D, S4E). Principal component analysis (PCA) separated the mutation carrier cell lines from controls in the first component (Figure S4F), indicating that expression variability between cell lines is mostly explained by the *CASK*-mutation status. The second component of the PCA separated samples by sex.

With support from the PCA results, we performed a pooled analysis of mutation carrier transcriptomes to detect consistent CASK-related expression changes. We detected 4098 differentially expressed transcripts with more than 20 reads at an adjusted p-value of 0.05 (Benjamin-Hochberg adjusted) and a 0.5 log2 fold change (LC) (Table S3). Gene set enrichment analysis (GSEA) revealed the upregulation of genes involved in morphogenesis and development of different tissues, as well as extracellular matrix components and signaling pathways WNT and BMP (Figure 4A). The downregulated categories consisted of presynaptic components, including the synaptic vesicle genes (Figure 4B). In addition, we were specifically interested in the enrichment of genes involved in protein-protein interactions (PPI) with CASK (Pathway Commons), genes linked to ASD (SFARI score 1, 2, and syndromic) and compiled list of NDD genes (Table S4). We detected significant enrichment of CASK PPIs in downregulated genes (p=0.003, FDR=0.003, GSEA, Figure 4C, Table S5) and no enrichment in upregulated genes (p=0.17, FDR=0.25, GSEA, Figure 4C, Table S5). Core enrichment genes are within a densely connected CASK PPI network, including MINT1 (encoded by *APBA1*), which is one of the tripartite binding partners of CASK (Figure 4D). We separately looked at the nuclear interaction partners TBR1, CINAP (*TSPYL2*), and BCL11A that were not included in the obtained CASK PPI dataset. We found downregulation of *TSPYL2* (LC=-0.37, p.adj.=2E-3, DESeq2) and *BCL11A* (LC=-0.53, p.adj.=1E-4, DESeq2), and absence of *TBR1* expression in our model. Moreover, we detected a nominally significant enrichment of NDD genes in upregulated genes (p=0.002, FDR= 0.25, GSEA, Figure 4C, Table S5). In addition, we performed GSEA enrichment for individual cases versus sex-matched controls (Figure 4C, Table S5-S7) and found that CASK PPI and NDD genes were more enriched in the pooled analysis, indicating that this approach allowed us to detect consistent expression changes, independent of genetic backgrounds. In a complementary approach, we overlapped the most significant DEGs of case versus sex-matched control analyses (Figure 4E). Over-representation processes confirmed extracellular structure organization (p<0.0001, FDR<0.0001, WebGestaltR, Table S8) and synaptic vesicle cycle genes (p<0.0001, FDR=0.0002, WebGestaltR, Table S9) as most significant up- and downregulated process, respectively. Results from pathway analysis demonstrated that *CASK*-mutant maturing neurons are deficient in expression of presynaptic CASK-interacting proteins.

**Figure 4.**
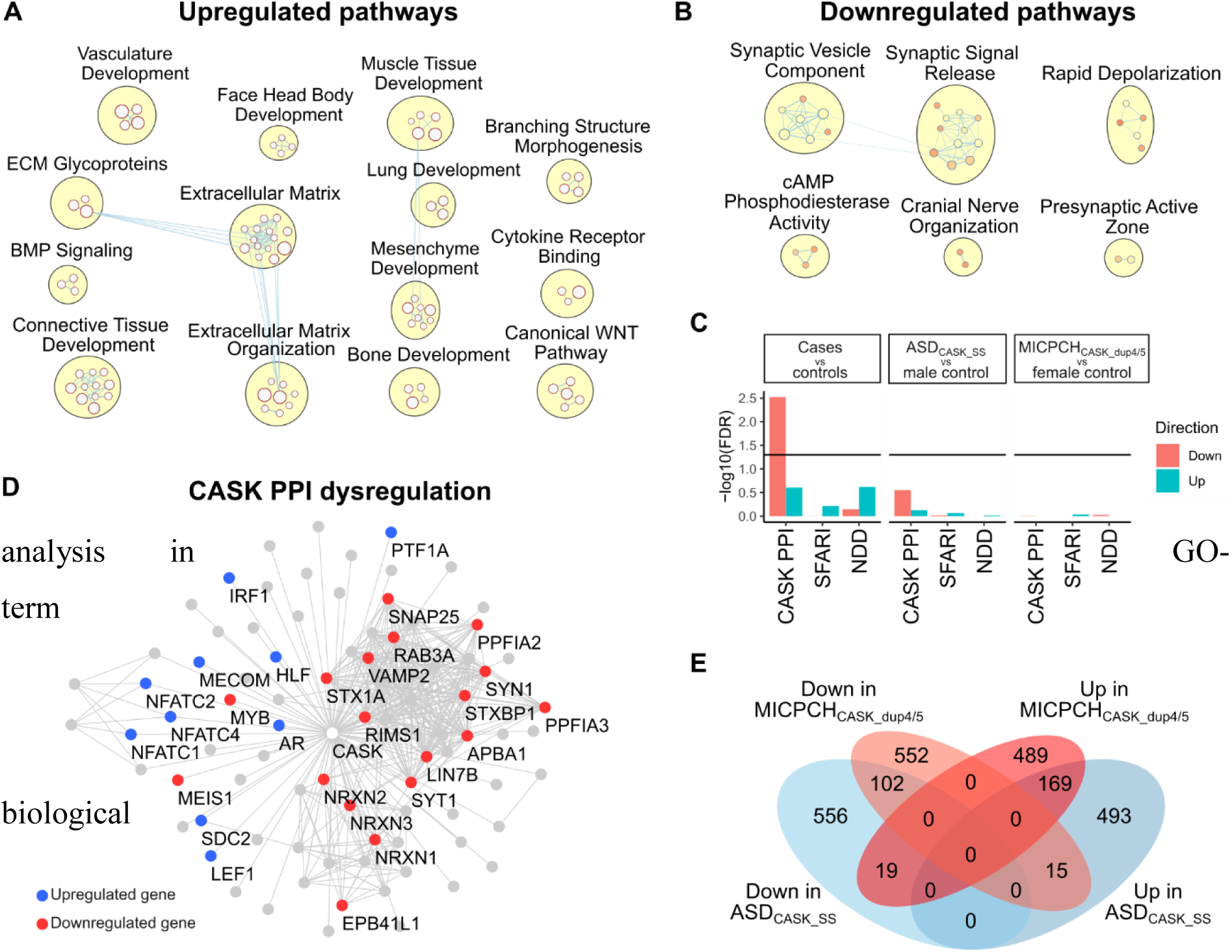
Consistent dysregulation of presynaptic and CASK interacting genes in bulk RNA-sequencing. (A) Upregulated and (B) downregulated pathways emerging from GSEA in bulk RNA-sequencing data from mutation carriers compared with controls. (C) GSEA for CASK interacting proteins (CASK PPI), ASD risk genes (SFARI) and NDD genes (NDD) in bulk RNA-sequencing data from mutation carriers compared with controls. (D) Network of CASK interacting proteins with upregulated marked in blue and downregulated in red. (E) Venn diagram illustrating top up- and downregulated genes in ASD_CASK_SS_ and MICPCH_CASK_dup4/5_ in comparison to sex-matched controls.

We performed siRNA-mediated knockdown of *CASK* to verify *CASK*-dependent expression changes during differentiation. We transfected male control cells after one-day of differentiation with a pool of CASK-specific siRNAs or non-targeting control siRNA. Knockdown was efficient with a significant reduction of CASK mRNA and protein at day 28 (Figure S4G, S4H). GSEA of DEGs confirmed changes in synaptic vesicle pathway and extracellular matrix compounds (Figure S4I). Similar to DEGs from *CASK-*mutation carrier cell lines, NDD genes were nominally enriched in upregulated genes after *CASK* KD (p=0.031, FDR=0.58, GSEA, Table S10). In contrast to reduced expression of synaptic vesicle genes in the *CASK*-mutation carrier neurons, the genes underlying synaptic vesicle enrichment in response to CASK KD were upregulated. The conflicting direction of expression changes may be due to priming effects of reduced CASK in NES cells before differentiation.

To test if the dysregulation of presynaptic genes results in aberrant presynaptic development, we assessed synapse morphology in differentiating neuronal cultures through immunofluorescence co-staining of pre- and postsynaptic markers Synapsin-1/2 and Homer-1, respectively (Figure 5A). Synapsin-1/2 is part of the CASK PPI network (Figure 4D), and the expression of SYN1 gene was downregulated (LC=-1.08, adj.p=3.03E-13, DESeq2). The Homer-1 gene, *HOMER1* also showed downregulation (LC=-0.33, adj.p=0.002, DESeq2). The number of Synapsin-1/2 positive presynaptic and Homer-1 positive postsynaptic regions did not reveal any difference between cell lines (Figure S5A). Similarly, the number of colocalizing regions, indicative of functional synapses did not differ. While the Homer-1 particle size was comparable between cell lines (Figure 5B), the mean Synapsin-1/2 particle size was smaller in ASD_CASK_SS_ neurons than controls (ASD_CASK_SS_ vs female control p=0.014; vs male control p=0.021, ANOVA posthoc Tukey, Figure 5C). Although dysregulation of presynaptic pathways was detected in the MICPCH_CASK_dup4/5_ cell line, the presynaptic Synapsin-1/2 size was comparable to controls. The size differences indicated that reduced *SYN1* expression affected presynaptic Synapsin-1/2 size in ASD_CASK_SS_. Differences in Synapsin-1 particle size and a more severe cellular phenotype in ASD_CASK_SS_ are in line with the results obtained for CASK particles.

**Figure 5.**
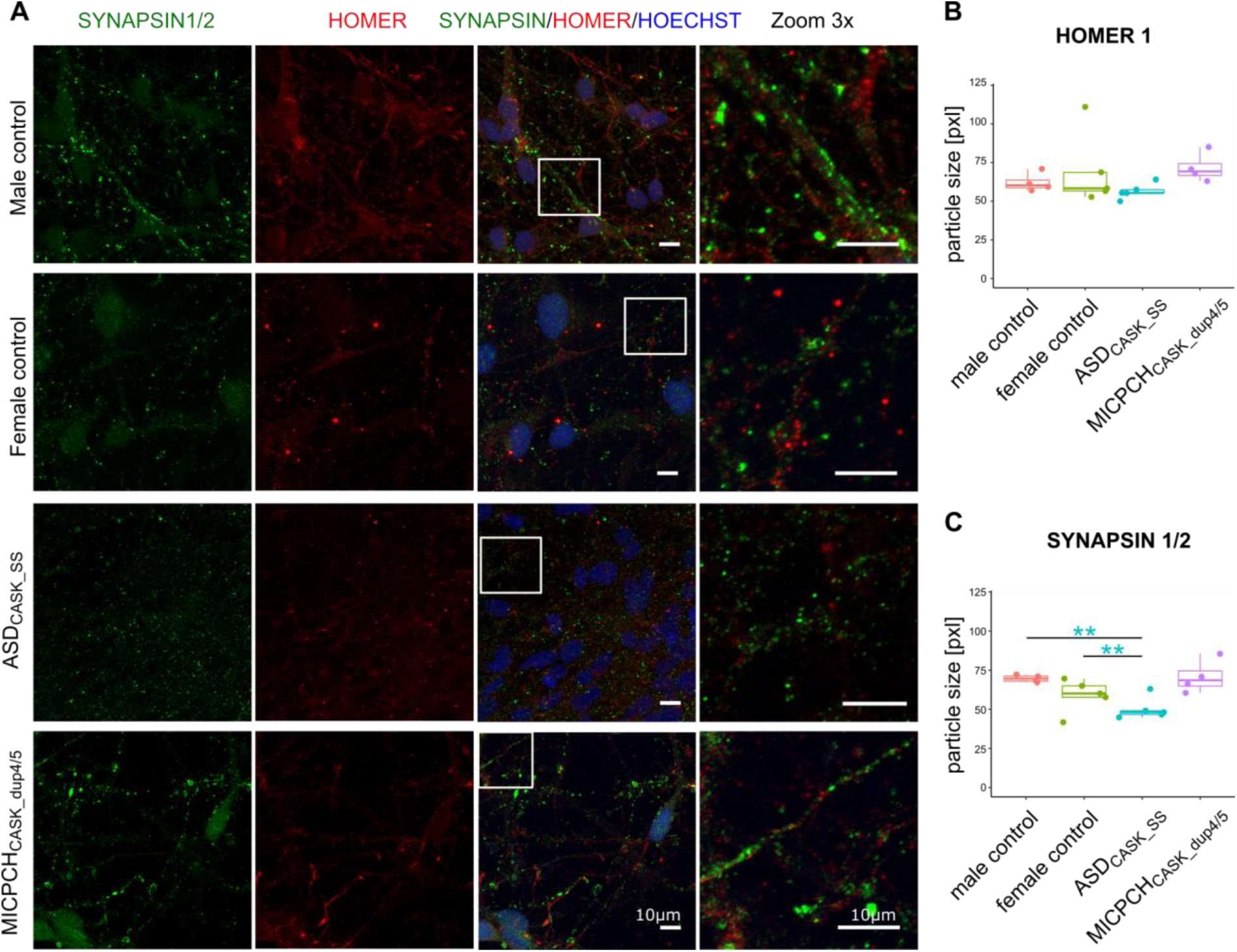
Reduced CASK levels decrease presynaptic Synapsin-1/2 marker size. (A) Representative confocal microscopy images of neurons differentiated for 28 days and immunostained for Synapsin-1/2 (green), Homer-1 (red) and Hoechst (blue). Scale = 10µm. (B) Quantification Homer-1 and (C) Synapsin-1/2 particle size. Female control (n=4), male control (n=4), ASD_CASK_SS_ (n=5) and MICPCH_CASK_dup4/5_ (n=4). Statistical differences between cell lines were calculated using ANOVA with post-hoc Tukey HSD. *p < 0.05, **p < 0.01, ***p < 0.001. Asterisks are colour-coded according to case cell lines.

### Excitatory-inhibitory balance is affected by reduced CASK levels

Next, we explored the effect of the *CASK* mutations on the identified cell types using the BSEQ-sc analysis pipeline for gene expression deconvolution (Baron et al., 2016). We used the cell type markers of the MICPCH_CASK_dup4/5_ case to predict cell type proportions underlying the bulk RNA transcriptomes. In our analysis, the estimated frequency of inhibitory GABA and undefined neuronal subtypes was unaffected in mutation carrier cell lines (Figure 6A). Excitatory neurons were predicted with a lower proportion in both case cell lines, and NPCs were predicted with a higher frequency in ASD_CASK_SS_. In accordance with changes in protein expression and morphological features, ASD_CASK_SS_ showed a stronger effect than MICPCH_CASK_dup4/5_. The deconvolution of the *CASK* KD transcriptome predicted no change in cell type proportions (Figure S5B).

**Figure 6.**
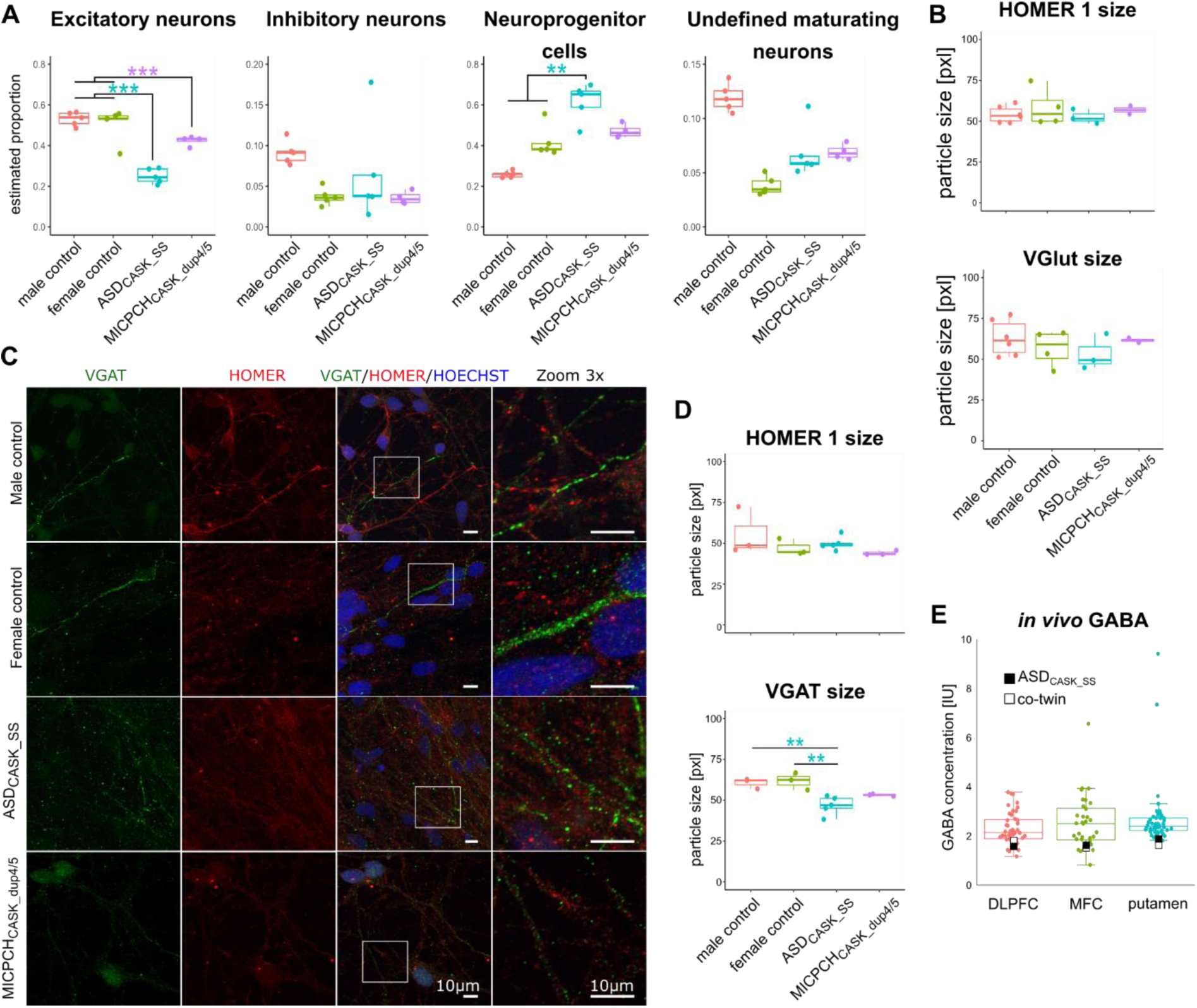
Presynaptic effect is limited to inhibitory VGAT presynaptic marker. (A) Deconvolution of cell type populations underlying bulk RNA-sequence samples from all cell lines. (B) Quantification of VGlut and Homer-1 particle size. Female control (n=4), male control (n=6), ASD_CASK_SS_ (n=3) and MICPCH_CASK_dup4/5_ (n=3). (C) Representative confocal microscopy images of neurons differentiated for 28 days and immunostained for inhibitory VGAT (green), Homer-1 (red) and Hoechst (blue). Scale = 10µm. (D) Quantification of VGAT and Homer-1 particle size. Female control (n=3), male control (n=3), ASD_CASK_SS_ (n=5) and MICPCH_CASK_dup4/5_ (n=3). Statistical differences between cell lines were calculated using ANOVA with post-hoc Tukey HSD. *p < 0.05, **p < 0.01, ***p < 0.001. Asterisks are colour-coded according to case cell lines. (E) *In vivo* concentration of GABA in the DLPFC, MFC and putamen of ASD_CASK_SS_ and his co-twin in relation to a typical developed control group.

Differences in cell type proportions should be detectable on a morphological level. We performed immunofluorescence staining of presynaptic markers VGlut and VGAT as a proxy for excitatory and inhibitory neurons, respectively. We had detected downregulation of *SLC17A6* encoding for VGlut (LC=-1.38, adj.p=2.44E-8, DESeq2) and *SLC32A1* encoding for VGAT (LC=-2.44, adj.p=1.09E-37, DESeq2) in the bulk RNA sequencing comparisons. VGlut particle number and size were comparable between cell lines (Figure 6B, Figure S5C). The consistent number of VGlut particles per nuclei indicated comparable numbers of excitatory neurons. Staining of the inhibitory marker VGAT showed significant decrease in particle size between ASD_CASK_SS_ and controls (ASD_CASK_SS_ vs female control p=0.005; vs male control p=0.009, ANOVA posthoc Tukey, Figure 6C and 6D). In addition, the size of colocalizing VGAT and Homer-1 particles, indicative of functional inhibitory synapses, was significantly reduced in ASD_CASK_SS_ (ASD_CASK_SS_ vs female control p=0.009; vs male control p=0.03, ANOVA posthoc Tukey, Figure S5D). The difference in colocalizing particle size was also detected between MICPCH_CASK_dup4/5_ and the sex-matched control (p=0.03, ANOVA posthoc Tukey). In agreement with previous findings for CASK and Synapsin-1/2 particles, the number of presynaptic particles was unaffected (Figure S5D). Reduced VGAT particle size indicates reduced inhibitory synapse size and suggests weaker inhibitory synapse strength. Comparable VGAT presynaptic particles per nuclei indicate that inhibitory neuron numbers are comparable.

To compare our iPSC-derived neuronal data of reduced VGAT staining in inhibitory synapses to *in vivo* neurotransmitter concentrations, we explored proton magnetic resonance spectroscopy ([1H]MRS) data, available for twin pairs including ASD_CASK_SS_ (Isaksson et al., 2018). We analyzed the GABA concentration in individuals without NDDs and obtained data for the dorsal medial prefrontal cortex (DLPFC, control n=44), medial frontal cortex (MFC, control n=33), and putamen (control n=45) (Figure S6A-D). The GABA concentration of the ASD_CASK_SS_ individual ranked in the lowest percentile in the DLPFC and Putamen and in the 9^th^ percentile of the MFC. GABA in the mutation carrier co-twin ranked similarly low in the DLPFC, with a slightly higher concentration (Figure 6E).

Finally, we investigated spontaneous neuronal activity in the neuronal cultures. We imaged intra-cellular calcium release in 4- and 5-week old neuronal cultures and compared calcium signaling events between the cell lines. We identified spontaneously active neurons in 4-week old cultures and observed an increased rate of calcium signals after 5 weeks (Figure 7A, B). No differences were detected between cell lines after 4 weeks of differentiation. At 5 weeks, ASD_CASK_SS_ and MICPCH_CASK_dup4/5_ showed less activity as compared with the male control (ASD_CASK_SS_ p=0.0266, MICPCH p=0.0019, Bonferroni corrected Pairwise Wilcoxon Rank Sum Test).

**Figure 7.**
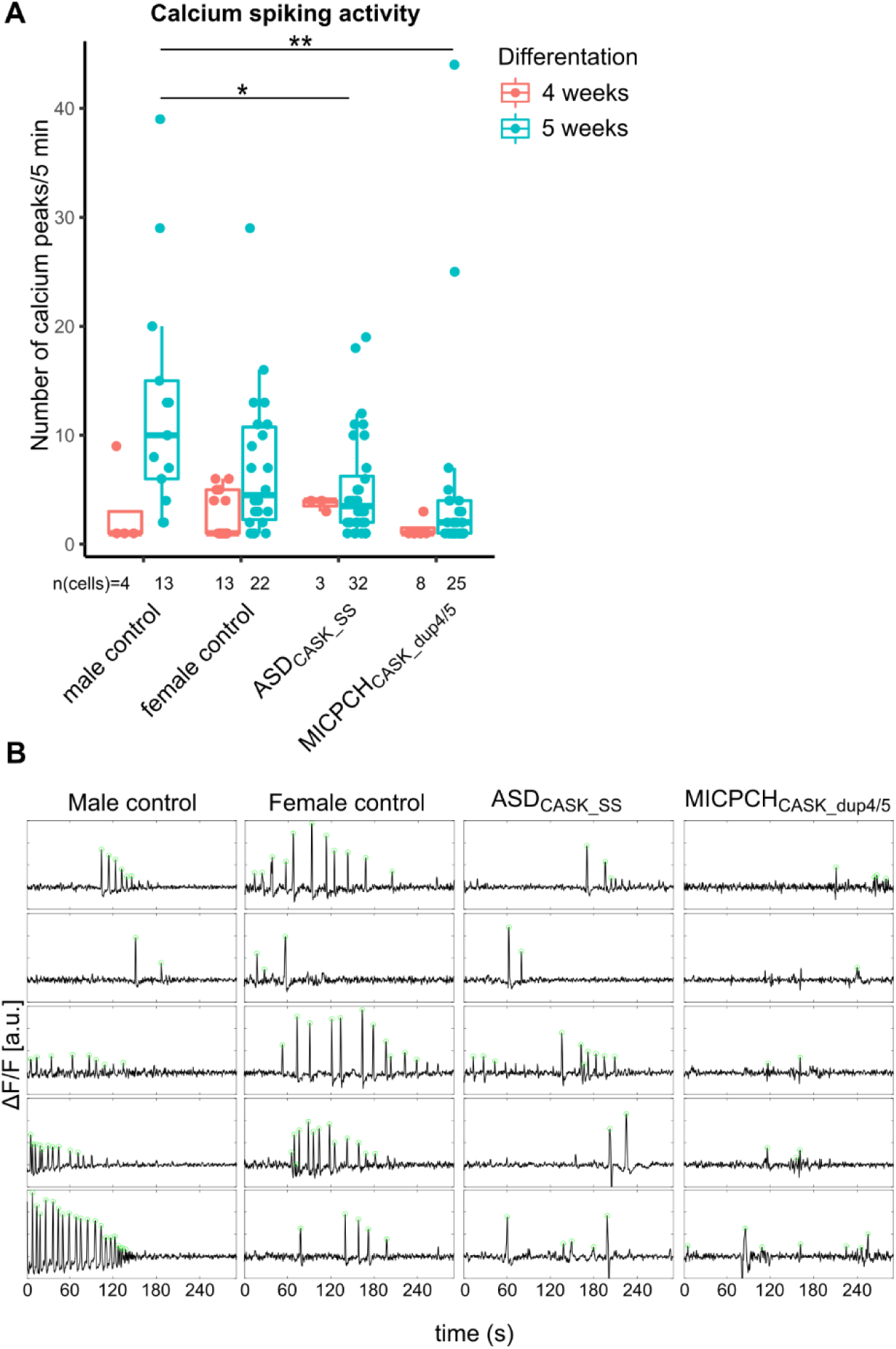
Neuronal activity measured by time-lapse calcium imaging. (A) Calcium peaks in active neurons of mutation carriers and controls differentiated for 4 and 5 weeks. Number of total neurons below x-axis. Biological replicates at 4-weeks: Female control (n=2), male control (n=2), ASD_CASK_SS_ (n=1) and MICPCH_CASK_dup4/5_ (n=2). 5-weeks: Female control (n=2), male control (n=2), ASD_CASK_SS_ (n=5) and MICPCH_CASK_dup4/5_ (n=2). (B) Representative five-minute traces of active neurons after 5-week differentiation with called peaks locations (green).

## Discussion

*CASK*-related disorders have emerged as an important genetic diagnosis for children with NDDs ranging from severely affected MICPCH patients to individuals with ASD (Deciphering Developmental Disorders, 2017; Hayashi et al., 2017; Iossifov et al., 2014; Moog et al., 2015). The role of CASK as an essential player in synapse formation was reported previously using model organisms (Atasoy et al., 2007; Butz et al., 1998; Mori et al., 2019); however, there is limited information of how the different *CASK* genetic variants found in humans affect the neuronal development. Therefore, we studied the consequences of two different mutations affecting *CASK* from individuals diagnosed with MICPCH and ASD. We utilized an iPSC derived model to investigate the mutation effects during early neuronal maturation with detailed molecular, cellular, and functional characterization. We show that even slight changes in CASK expression in humans lead to dysregulation of the formation of presynapses, especially in inhibitory neurons. We suggest that the revealed presynaptic phenotype could be underlying the social difficulties seen in individuals affected by *CASK*-related disorders and that there are other mechanisms leading to the more profound brain malformations seen in more severely affected patients.

Using transcriptomic profiling of maturing neurons from *CASK* mutation carriers and controls, we demonstrated that reduced CASK levels affected the transcription of presynaptic genes with significant enrichment for CASK-interacting partners. Importantly, downregulation of, e.g. *NRXN*-family of genes, *STXBP1,* and *SYT1* in *CASK* mutation carriers provide a phenotypic link to other NDD genes. For instance, *NRXN1* is one of the most studied NDD genes in which homozygous mutations cause Pitt-Hopkins-Like syndrome 2 characterized by NDDs such as ID and ASD (Zweier et al., 2009) and heterozygous deletions predispose to several NDDs, psychiatric disorders as well as congenital malformations (Al Shehhi et al., 2019; Dabell et al., 2013). Mutations in *STXBP1* were first described in patients with early infantile epileptic encephalopathy and are a common cause of epilepsy and encephalopathy with ID (Saitsu et al., 2008; Stamberger et al., 2016; Uddin et al., 2017). *SYT1* mutations in Baker-Gordon syndrome cause infantile hypotonia, ophthalamic abnormalities, hyperkinetic movement disorders, motor stereotypies and developmental delay (Baker et al., 2018). Disruption of this protein network in the developing presynapse could be underlying the behavioral phenotypes underlying *CASK*-related NDDs. In addition to the presynapse, CASK has been implicated in functions within the dendrites, postsynapse, and neuronal nucleus. Although we observed the strongest enrichment in the presynapse, functional interaction partners of CASK in the other neuronal compartments were also dysregulated. We observed upregulation of *CNTNAP2* and *SDC2*, which form complexes with CASK in dendrites to regulate postsynaptic development (Gao et al., 2018; Gao et al., 2019; Hsueh and Sheng, 1999; Hu et al., 2016). *CNTNAP2* is linked to Pitt-Hopins-Like syndrome 1 and increased ASD risk (Bakkaloglu et al., 2008; O’Roak et al., 2011; Zweier et al., 2009). In addition, we observed downregulation of nuclear interacting partners *BCL11A* and *TSPYL2*, which have been linked to ID and ASD, respectively (Dias et al., 2016; Moey et al., 2016). Taken together, *CASK* mutations cause dysregulation of NDD-related networks with a strong effect on presynapse development and putative impact on additional aspects of neuronal functions. Interestingly, we also illustrate upregulation of many developmental pathways in our analyses such as mesenchyme, face and head development as well as WNT signaling. Future studies should investigate these pathways in more detail and characterize how they might be linked to *CASK*-related phenotypes, including microcephaly, scoliosis, and abnormalities of the eye and skin development (Deciphering Developmental Disorders, 2017).

One benefit of the neuronal differentiation scheme used here was that we could investigate the effects on both inhibitory and excitatory neurons, and demonstrate that the transcriptional changes translate to reduced presynaptic size of inhibitory neurons as measured by Synapsin-1/2 and the inhibitory marker VGAT. The reduced presynaptic size, as measured by the selected markers and CASK aggregates, may indicate aberrant synaptogenesis and decreased synaptic strength. In turn, this could lead to E/I imbalance. Earlier studies using electrophysiological measures in transgenic mice demonstrated decreased frequencies of mIPSCs and increased mEPSCs in adult CASK-deficient neurons (Atasoy et al., 2007; Mori et al., 2019), establishing aberrant E/I balance as a phenotype underlying CASK-deficiency. In addition, Mori et al. (2019) detected a decreased expression of the glutamate receptor GluN2B (encoded by *Grin2b*) in CASK-deficient neurons and proposed that reduced postsynaptic excitability caused an increase in excitatory output as a compensatory mechanism. In line with the previous findings, we detected reduced *GRIN2B* expression in ASD_CASK_SS_ during neuronal maturation. However, we detected an unaffected number and size of excitatory VGlut synapses in contrast to the reduction in inhibitory presynaptic size, which together suggest that reduced inhibitory signaling is the primary mechanism underlying E/I imbalance in CASK-deficient neurons. Moreover, the low GABA concentration in all assessed brain regions of ASD_CASK_SS_ and his co-twin could indicate that the reduced inhibitory signal strength is persistent during postnatal development. On the *in vitro* level, we observed reduced neuronal activity in the mutation carrier neurons. The observed decrease in VGAT at GABAergic synapses could explain reduced bursting frequency. While a reduced inhibitory frequency would be expected to increase network activity, reduced bursting complexity could explain the reduced number of neuronal calcium events. In addition, GABA neurotransmitter function changes from excitatory to inhibitory during neural development and may have had an excitatory function in our model system (Ben-Ari, 2002). Electrophysiological characterization of mISPCs and mEPSCs in ASD_CASK_SS_ and MICPCH_CASK_dup4/5_ neurons during neuronal maturation may further improve our understanding of primary and compensatory mechanisms underlying the E/I imbalance in CASK-related NDDs. E/I imbalance caused by aberrant neuronal firing was described for a variety of genetic mutations underlying NDD phenotypes. Studies investigating *NRXN1* showed that similar to *Cask* KO mice, *Nrxn1* KO mouse neurons remain with normal postsynaptic current amplitudes, but in contrast to *Cask* KO showed decreased mEPSC frequencies (Etherton et al., 2009). In human embryonic stem cell-derived neurons with heterozygous *NRXN1* LoF mutation, increased CASK protein levels were observed together with decreased mEPSC frequencies (Pak et al., 2015) and homozygous deletions impair neuron maturation (Lam et al., 2019), indicating that CASK-related pathways affect E/I balance. In a recent study of isogenic human excitatory neurons disrupted for ASD-relevant genes, electrophysiological assessment demonstrated reduced spontaneous EPSCs and neuronal bursting in the presence of homo- or hemizygous mutations of *ATRX*, *AFF2*, *KCNC2*, *SCN2A* and *ASTN2* (Deneault et al., 2018). Another study detected reduced neuronal firing in iPSC-derived neurons of eight individuals with idiopathic ASD (Marchetto et al., 2017) and detected a correlation of reduced network complexity to behavioral and cognitive phenotype of the donors (Amatya et al., 2019). Moreover, reduced GABA concentration, as seen in the ASD_CASK_SS_ brain was reported in groups of children with ASD (Ford and Crewther, 2016). Taken together, these findings show that *CASK*-related NDDs share the E/I imbalance as pathological mechanisms with other ASD- and NDD-risk genes.

In addition to the effects seen overall for the neuronal cultures, our study shows intriguing findings from the two unique mutations affecting *CASK*. The hemizygous splice-site mutation detected in an autistic individual and his mild symptomatic monozygotic co-twin reduced expression of CASK_WT_ and caused splicing of two mutant mRNA isoforms. We excluded gain-of-function effects from mutant mRNA isoforms, as we detected only wild type protein with similar cellular localization as in matched control neurons. Interestingly, we observed the rescue of the *CASK*_WT_ splicing by an unknown mechanism. RNA sequencing of ASD_CASK_SS_ neurons indicated that the mutated T/U is modified to the reference G nucleotide, suggesting that RNA editing is involved in the rescue mechanism. While reports on U-to-G RNA editing are rare, these mechanisms are shown to be relevant for synaptic transcripts and may be involved in the rescue of *CASK*_WT_ in the ASD_CASK_SS_ (Dai et al., 2018; Reid et al., 2008). As both monozygotic twins carry the splice site mutation and exhibit differences in symptom severity, it is interesting to speculate that environmental stressors could influence the rescue mechanisms and thus lead to variable severity. We have earlier studied pre- and perinatal environmental exposures in the RATSS twin sample, including the ASD_CASK_SS_ case and his co-twin, and showed that uptake of essential metals zinc and manganese, and the neurotoxic metal lead were associated with variable manifestation of autistic traits and ASD diagnoses (Arora et al., 2017). These environmental stressors could potentially affect the RNA processing efficiency and mediate gene-environment interactions leading to differences in outcomes of the twin pair (Dai et al., 2018; Dick et al., 2019; Shomron et al., 2002).

The instability of XCI has been reported in iPSCs and has hindered the cellular investigation of dominant X-linked genes in NDDs. In the MICPCH_CASK_dup4/5_ cells of the female donor, we observed mutant and wild type CASK mRNA expression that likely resulted from XCI escape of the wild type allele. Reports of XCI in adult human tissues did not indicate CASK as likely XCI escape gene (Tukiainen et al., 2017). Heterozygous CASK knockout mice that recapitulate the MICPCH phenotype showed *Cask*_WT_ expression in 50% of neurons suggesting that XCI is stable in *CASK*-negative neurons (Mori et al., 2019; Srivastava et al., 2016). The stability of XCI during iPSC re-programing varies between protocols, and activation of both X chromosomes with subsequent random silencing of one X has been observed (Dandulakis et al., 2016). The milder phenotype in females carrying *CASK* mutations with skewed XCI suggests that the proportion of *CASK*-deficient cells is crucial for phenotype severity (Moog et al., 2011; Seto et al., 2017). Thus, the escape of *CASK*_WT_ in the MICPCH_CASK_dup4/5_ cells is unlikely to recapitulate all pathoetiological mechanisms in cases with MICPCH. Indeed, we consistently observed less severe cellular phenotypes in the MICPCH_CASK_dup4/5_ cells than ASD_CASK_SS_ cells. This demonstrates the necessity to control for XCI pattern in iPSC studies of X-linked genes.

Currently, there are no drugs to treat *CASK*-related disorders. In a recent clinical study assessing neurorehabilitation effects on three girls with MICPCH and *CASK* mutations at the age of 2 to 4 years showed that the girls improved in motor skills, social interaction and communication functions, indicating potential benefits of targeting the social aspects in *CASK-r*elated disorders (DeLuca et al., 2017). As we demonstrate a specific effect on the inhibitory presynapse and reduced GABA concentration in the cortical and subcortical brain regions of ASD_CASK_SS_, it is intriguing to speculate on the effect of GABA modulators in *CASK*-related disorders. Especially GABA agonists could be tested, which have shown positive effects on social deficits in ASD and are in clinical trials for Fragile-X syndrome (Erickson et al., 2017; Erickson et al., 2014a; Erickson et al., 2014b). Further development of drugs that target the presynapse and synaptic vesicle cycle could also be beneficial but are currently not well explored (Li and Kavalali, 2017).

In summary, we show that reduced CASK protein levels affect presynaptic development and decrease inhibitory synapse size, which may alter E/I balance in developing neural circuitries. Aberrant E/I balance, and synaptogenesis are two common biological pathways underlying NDDs of different genetic origin. Future pharmacological and clinical studies on targeting presynapses and E/I imbalance could lead to specific treatments for *CASK*-related disorders.

## Materials and Methods

### Identification of mutation carriers

The mutation carriers were recruited to the study from two different sources: The Roots of Autism and ADHD study in Sweden (RATSS) and Clinical Genetics at the Karolinska University Hospital. The comprehensive phenotyping procedures for the twin pair has been described elsewhere (Bölte et al., 2014; Myers et al., 2017) and included T1-weighted spoiled gradient echo anatomical MRI scans (176 slices, TR=8.2s, FOV=240; acquired with a 3 Tesla MR750 GE scanner). Phenotyping, genotyping and clinical assessment of the girl with MICPCH was described earlier (Wincent et al., 2015). Written informed consent was obtained from affected individuals and their parents prior to the study. The study was approved by the regional and national ethical boards in Sweden, and it has been conducted in accordance with the Declaration of Helsinki for medical research involving human subjects, including research on identifiable human material and data.

### Whole-exome sequencing and data processing

For the identification of the genetic alterations, we performed both microarray analysis (for both cases) and whole exome sequencing (for the twin pair and their parents). For *de novo* variant detection, the parents were included in the genetic analyses. DNA was extracted from whole blood and saliva samples using standard methods. The detection of CNVs has been described earlier (Stamouli et al., 2018; Wincent et al., 2015). For WES, 3 μg DNA (200 ng for the mother) was used for library preparation with Agilent SureSelect Human All Exon V5 kit followed with quantification of the libraries using KAPA Biosystems’ Library Quantification according to the manufacturer’s recommendations. The sequencing was performed on Illumina HiSeq 2500 using the Rapid SBS Kit v2 chemistry. After sequencing, base calling was done with Illumina Casava pipeline v1.8.2, and reads were mapped to the hg19 reference sequence using the Burrows-Wheeler Aligner (BWA) v0.5.9. Duplicate reads were marked and removed using Picard v1.79 tools. Variant calling, local realignment around indels and base quality score recalibration were conducted with Genome Analysis ToolKit (GATK) v1.1-28. Basic variant calling was followed by annotation using a custom pipeline at the Center of Applied Genomics (TCAG) based on Annovar that creates a merged file for the twin pair and parents (Tammimies et al., 2015). Population frequency for each variant was based on the 1000 Genomes Project, the Exome Variant Server hosted by NHLBI (Exome variant server), 69 Complete Genomics public genomes and The Exome Aggregation Consortium (ExAC). Additional information about putative pathogenicity and disease association of variants were based on different databases such as dbSNP, Clinvar, Online Mendelian Inheritance of Man (OMIM), and Clinical Genomics Database (CGD). Reported mutation sites were transferred to hg38 using LiftOver (UCSC Genome Browser).

### Identification of putative risk variants underlying the NDD phenotype

For the analysis of putative NDD risk variants seen in both ASD_CASK_SS_ and his co-twin, only rare variants (< 1.0% in the population) were selected. The remaining variants were filtered based on the impact of the variant and association of the gene to NDDs. In brief, the genes affected by the prioritized variants were compared against a list of known ASD and ID genes (Pinto et al., 2014) and genes implicated in neurodevelopmental/behavioral phenotypes in human or mouse from the human phenotype ontology (HPO) (Kohler et al., 2014). Additionally, a list of genes implicated in ASD from two large scale sequencing studies (De Rubeis et al., 2014; Iossifov et al., 2014) was compiled and used in the analysis. The mode of inheritance for the genes was curated from the OMIM and CGD databases. Further prioritization was made based on the predicted pathogenicity of the variants; *de novo* variant, LoF (loss of function, i.e. stop-gain, frameshift, essential splice site alterations, or missense near splice-site) and missense variants were selected for further evaluation. Final categorization was made based on the criteria from the American College of Medical Genetics and Genomics (ACMG) (Richards et al., 2015).

To validate genomic variants, genomic regions were PCR amplified using sequence specific primers with HotStarTaq Polymerase Kit (Qiagen # 203203). PCR primer sequences used to amplify the CASK_SS variant: AGAAAATCCCTTGTCTGATGA (Fw) and CCTGCCATAAAAATCCACTC (Rv). DIAPH1 variant: TTACTTACCCCCAACACAAAC (Fw) and TTACTGAGATGGAGAGCTTGC (Rv). The PCR products were purified, and Sanger sequenced.

### Generation of neuroepithelial stem cells

Human iPSC cells were derived for the *CASK* mutation carriers diagnosed with MICPCH and ASD, using a previously described procedure (Uhlin et al., 2017). In summary, fibroblasts were obtained from skin biopsies and reprogrammed to iPSC using Sedai virus transfection of the Yamanaka factors *OCT4*, *SOX2*, *KLF4,* and *cMYC*. Human iPS cells were maintained in xenofree and defined conditions using, substrate Laminin521 (BioLamina #LN521), and Essential E8 media (ThermoFisher Scientific # A1517001). Quality of iPSC induction was done with NANOG and OCT4 staining (Figure S6A) and gene-expression based pluripotency test (Muller et al., 2011). Genetic identity was controlled with karyotyping (Figure S6B) and short-tandem repeat analysis of 20 markers.

iPS cells were passaged as single cells by TrypLE Select (1X) (Gibco # 12563029) once they reached 80-90% confluence. Dual-SMAD inhibition was applied to derive NES cells from human iPS cells as described in Chambers SM et al. 2009 (Chambers et al., 2009). On the first day of neural induction, iPS cells were seeded at a density of 40 000 cells/cm^2^ on LN521 substrate and in Essential E8 medium. Day 1-5 of Neural Induction, E8 medium was replaced with KRS-medium including the human recombinant Noggin (hNoggin, PeproTech #120-10C) at a final concentration of 500ng/ml, SB431542 (StemCell Technologies #72232) at a final concentration of 10 μM and CHIR99021 (StemCell Technologies #72052) at a final concentration of 3,3 μM. The media was replaced daily. Day 6-12, KOSR media was gradually replaced increasing amounts of N2-media (25%, 50%, 75%), according to Chambers at al 2009, maintaining the factors hNoggin at a final concentration of 500 ng/ml and CHIR99021 at a final concentration of 3,3 μM. At day 12 of neural induction, cells were dissociated by TrypLE Express, and seeded in DMEM/F12+Glutamax medium (Gibco) supplemented with 0.025X B-27 (Gibco), 1X N-2 (Gibco), 10 ng/ml recombinant human FGF (Gibco), 10 ng/ml recombinant human EGF (Peprotech) and 10 U/ml Penicillin/Streptomycin (Gibco). Cells were seeded at a density of 250 000-280 000 cells/cm^2^, on plastic surface double-coated with 20 ug/ml poly-ornithine (Sigma Aldrich) and 1 ug/ml laminin (Sigma Aldrich). The cells were further passaged at a density of 40 000 cells/cm^2^. Half of the medium was changed daily.

### Differentiation of NES cells

NES cells were seeded on plastic surface, pre-coated with 20 ug/ml poly-ornithine (Sigma Aldrich) and 1 ug/ml laminin (Sigma Aldrich) in DMEM/F12+glutamax medium (Gibco) supplemented with 0.5X B-27 (Gibco), 1X N-2 (Gibco) and 10 U/ml Penicillin/Streptomycin (Gibco). For immunofluorescence, neurons were seeded on a glass surface, pre-coated with 200 ug/ml poly-ornithine (Sigma-Aldrich), and 4 ug/ml laminin (Sigma-Aldrich). Cells were maintained in a 5% CO_2_ atmosphere at 37°C. Two third of the medium was changed every second day of differentiation with 0.4 ug/ml laminin added. Neurons were differentiated for 8, 16 or 28 days. Cells harvested for time point day 0 were cultured for two days in NES cell medium.

### RNA extraction and RT-qPCR

Cells were lysed in TRIzol reagent (Invitrogen), and RNA was isolated using the ReliaPrep RNA Cell Miniprep (Promega #A1222). The RNA was reverse transcribed using iScript cDNA Synthesis Kit (BioRad) and cDNA quantified with SsoAdvanced Universal SYBR Green Supermix (BioRad) following manufacturer protocols on a CFX96 thermal cycler (BioRad). CFX Manager software was used to record amplification curves and to determine Ct values. RT-qPCR reactions were performed in technical triplicates. We calculated the ΔCt to the GAPDH housekeeping gene and ^ΔΔ^Ct to control cell lines. We used three biological replicates obtained in individual experiments, if not stated otherwise in the figure legends. Statistical significance between cell lines was determined with ANOVA and posthoc Tukey HSD in R.

### Immunodetection using a capillary western blot

Cells were dissociated in extraction buffer (50 mM Tris-HCl, 100 mM NaCl, 5 mM EDTA, and 1 mM EGTA) using plastic cell scraper and lysed with 6 short sonication bursts at 36% amplitude (Vibra-Cell VCX-600, Sonics). Total protein yield was quantified with Pierce BCA assay (Thermo Fisher), and 250 µg/ml protein was loaded for Simple-Western WES (ProteinSimple) quantification. Antibodies were used to detect the proteins CASK (1:500 Novus Biologicals NBP2-41181), beta-actin (1:100 Abcam ab8227), and GAPDH (1:5000 Sigma G9545). The Compass Software For Simple Western (Version 4.0.0) was used to identified peaks at known molecular weights. In the chromatogram, the peak area was used for protein quantification. We normalized CASK protein for housekeeping protein. For time point days 0, 8, and 16, we obtained two biological replicates and three biological replicates for time point 28 and siRNA-mediated knockdown of CASK. Statistical significance between cell lines was determined with ANOVA and posthoc Tukey HSD in R.

### Immunofluorescence

Cells were cultured on glass coverslips for the indicated differentiation time. They were fixed for 20 min in 4% formaldehyde, rinsed with 1X TBS and blocked with blocking buffer (1X TBS, 5% Donkey Serum, 0.1% Triton X-100) for 1h. The slides were incubated with the primary antibodies diluted in blocking buffer overnight at 4°C. Primary antibodies used are CASK-NBP2-41181, 1:500 (Novus bio), MAP2-M2320, 1:500 (Sigma Aldrich), VGLUT1-135304, 1:250 (Synaptic system), Homer-1-160011, 1:250 (Synaptic system), VGAT-131003, 1:500 (Synaptic system), Synapsin-1/2-106006, 1:500 (Synaptic system), Nestin-MAB5326-KC, 1:1000 (Merk-Millipore), SOX2-AB5603, 1:1000 (Merk-Millipore). Slides were thoroughly washed with 1X TBS and incubated for 2 hours with secondary antibodies, and 1:500 Hoechst diluted in blocking buffer. The coverslips were washed and mounted with Diamond Antifade Mountant (Thermo Scientific). All images were taken with LSM 700 Zeiss Confocal Microscope, with 63x magnification at 1024 x 1024 pixel [pxl] resolution, resulting in an aspect ratio of 0.099233 µm per pixel.

CASK particles, as well as nuclei, were counted with the ImageJ Particle Analyzer, after converting fluorescent images to binary images (threshold function “Moments” for CASK and “Default” for nuclei). Quantification of particle number and size was done in R. Two independent replicates were obtained for the female control, three for MICPCH_CASK_dup4/5_ and four for ASD_CASK_SS_ and the male control. Of each independent replicate, we obtained two images (technical replicates). Statistical differences between cell lines were calculated on independent replicates, using ANOVA with posthoc Tukey HSD.

Synaptic marker particle size and number were quantified with the ImageJ plugin Synapse Counter, using default settings (Dzyubenko et al., 2016). For the Synapsin-1/2 – Homer-1 co-staining, we obtained four independent replicates for the female control, male control and MICPCH_CASK_dup4/5_, and five for ASD_CASK_SS_. For the VGlut – Homer-1 co-staining we obtained four independent replicates for the female control, six for the male control and three for MICPCH_CASK_dup4/5_, and ASD_CASK_SS_. For the VGAT – Homer-1 co-staining we obtained three independent replicates for the female control, male control and MICPCH_CASK_dup4/5_, and five for ASD_CASK_SS_. We obtained one to three images (technical replicates) per independent replicate. Statistical differences were calculated for mean particle size and number on independent replicates, using ANOVA with posthoc Tukey HSD.

### siRNA-mediated gene knockout

NES cells were seeded for differentiation and the next day transfected with 0.5µM Accell SMARTpool siRNA targeting *CASK* mRNA (Dharmacon #E-005311-00-0010) or 0.5µM Accell Green Non-targeting siRNA (Dharmacon #D-001950-01-20) according to standard protocol. Three biological replicates were harvested for each protein and RNA analysis.

### Single-cell RNA sequencing

Cells were dissociated from culture surface with 2 min trypsin incubation and subsequent trypsin inhibition. After 3 minutes’ centrifugation at 300 rcf the cells were resuspended in cold Dulbecco’s PBS and single cells sorted by size into lysis-buffer using BD FACS Aria III. Smart-Seq2 library preparation and sequencing were done with the Eukaryotic Single Cell Genomics (ESCG) facility in the SciLifeLab, Stockholm (Picelli et al., 2013). 384 sorted wells were sequenced with a total of 209.6 M reads and an average sequence depth of 550,000 reads per cell. Sequence reads were demultiplexed using deindexer and aligned to reference genome GRCh38/hg38 (STAR). Approximately 80% of uniquely aligned reads aligned to the genome, 40% to exons and 30% to introns (Figure S3A-B). The individual counts per gene and cell were reported in a count matrix (GSE140572) and used for further analysis. The scImpute package was used to calculate dropout expression values from the count matrix (Li and Li, 2018). Cells with less than 50,000 read counts in less than 2,000 genes were removed from analysis. Differential gene expression and cell clustering were done using the Seurat package (Butler et al., 2018). To obtain mutant CASK reads, we created a custom genome with STAR that included the cDNA sequence of the CASK gene with the duplicated exons 4 and 5 and re-aligned all sequence reads. Exon 5 to exon 4 spanning sequence reads were analyzed using vcftools.

### Bulk RNA sequencing

We extracted RNA samples of five biological replicates per cell line that were obtained in independent experiments. Samples were delivered to NGI Sweden for library preparation and sequencing. In short, strand-specific RNA libraries were prepared using Tru-Seq, including ribosomal depletion (Illumina). Samples were sequenced on NovaSeq6000 (NovaSeq Control Software 1.4.0/RTA v3.3.3) with a 2×151 setup using ‘S1’ flow cell mode. The Bcl to FastQ conversion was performed using bcl2fastq_v2.19.1.403 from the CASAVA software suite. The quality scale used is Sanger / phred33 / Illumina 1.8+. The NGI-RNASeq pipeline was used to trim reads (TrimGalore), align to reference genome GRCh38/hg38 (STAR), perform quality control (FastQC, RSeQC, dupRadar, MultiQC) and count gene reads (featureCounts). We obtained on average 50.4 million reads per sample with a minimum of 96.9% reads aligned to protein-coding regions. Sample read counts are supplied in GSE140572. Replicates of each cell line cluster together with the exception of one MICPCH replicate that was removed from analysis. Read alignment at the CASK gene locus were visualized in the Integrative Genomics Viewer (IGV v2.5.3).

Differential gene expression was calculated from gene counts using DESeq2 (v1.24.0) in R. To determine a differential expression for individual cases, we used passage and cell lines as independent variables and compared ASD_CASK_SS_ with male control and MICPCH_CASK_dup4/5_ with female control. Top ranking genes for comparison between ASD_CASK_SS_ and MICPCH_CASK_dup4/5_ were filtered at an adjusted p-value of 1E-5 (Benjamin-Hochberg adjusted), with more than 20 reads and log2 fold change (LC) higher than 0.5. The Venn diagram was visualized in R using the VennDiagramm package (v1.6.20), and over-representation analysis was performed using WebGestaltR (v0.4.2).

### Curation of NDD gene list

A list of known ASD and NDD genes were obtained from three online databases: Simons Foundation Autism Research Initiative (SFARI, https://gene.sfari.org/), DatabasE of genomiC varIation and Phenotype in Humans using Ensembl Resources (DECIPHER, https://decipher.sanger.ac.uk/ddd#overview), and Human Phenotype Ontology (HPO, https://hpo.jax.org/app/). All genes from DECIPHER, category 1, 2, and syndromic genes from SFARI and behavioral abnormality (HP:0000708) and cognitive impairment (HP:0100543) genes from HPO were included in the NDD gene list. Additionally, we included genes implicated in ASD, intellectual disability (ID) and developmental delay (DD) from five recent whole genome/exome sequencing and chromosome microarray studies (Coe et al., 2019; Pinto et al., 2014; Sanders et al., 2015; Satterstrom et al., 2019; Yuen et al., 2017). The date of data access is included in Table S4.

### Gene Set Enrichment Analysis

The unfiltered gene expression list was ranked by fold-change expression changes. Gene set enrichment analysis was performed using the GSEA desktop software (Version 3.0) (Subramanian et al., 2005). We used weighted enrichment statistics and included pathway gene set sizes between 3 and 250 genes. Pathway gene sets were obtained from Reimand et al. (Reimand et al., 2019). Enriched categories were visualized in Cytoscape (v3.7.0) with Enrichment Map (v3.1.0) and annotated with AutoAnnotate (Merico et al., 2010; Shannon et al., 2003).

Specific GSEA was performed on CASK PPI (downloaded from PathwayCommons; PCViz: CASK on 08.Oct.2019), the NDD gene list, and SFARI sub-list. Specific GSEA was performed on ranked gene lists split into up- and downregulated genes. The CASK PPI network was visualized in Cytoscape.

### Deconvolution

The deconvolution of bulk RNASeq data was done using the BSEQsc package (Baron et al., 2016). BSEQ-sc uses cell type specific marker genes from single-cell RNA transcriptomes to predict cell type proportions underlying bulk RNA transcriptomes. Deconvolution was done on increasing numbers of significant genes, and the predictions were stable when using 10 or 20 most significant marker genes. Statistical differences in the estimated cell proportions between patient and both control cell lines were done using ANOVA and posthoc Tukey HSD test.

### Calcium Imaging

Calcium events from neuronal activity last for 0.5 to 1 s and may be caused by individual action potentials or bursts of several action potentials (Forsberg et al., 2017; Pachitariu et al., 2018). Intracellular calcium recordings were performed using a Fluo-8 AM dye (AAT Bioquest). A dye stock was prepared containing 0.3 mM Fluo-8 AM and 3.3% Pluronic F-127 (ThermoFisher Scientific) and mixed thoroughly. Stock was added to the cell culture media to a final dilution of 5 µM Fluo-8 and 0.05% Pluronic F-127. The medium was carefully exchanged, and cells were incubated for 45 min at 37°C.

Then culture dishes were transferred to an upright Zeiss Examiner.D1 microscope with a pE-300 light source (CoolLED) and a prime 95B camera (Photometrics). After an adjustment period of 5 minutes, time-lapse videos (20Hz, 40ms exposure, 5 min) were recorded in two random spots on the dish using a Zeiss 40x water immersion objective with an NA of 1. Recordings were performed using a windows computer running Micro-Manager (Open Imaging) (Edelstein et al., 2014).

Image time series were pre-processed to balance signal loss from fluorescence decay using the ΔF/F method in ImageJ (Jia et al., 2011). Regions of interest (ROI) at active neurons with high variability of calcium signal were manually identified. Mean gray values of ROIs were recorded over the 5-minute pre-processed time series to obtain traces of neuronal activity. Traces were resampled to 2 Hz and smoothed using the python scipy.signal package. Peaks were detected using the PeakCaller Matlab software (Artimovich et al., 2017) with the following settings: 25% required rise, 25pts max lookback, 25% required fall, 25pts lookahead, Finite Difference Diffusion, 60 trend smoothness. Biological replicates at 4-weeks: Female control (n=2), male control (n=2), ASD_CASK_SS_ (n=1) and MICPCH_CASK_dup4/5_ (n=2). 5-weeks: Female control (n=2), male control (n=2), ASD_CASK_SS_ (n=5) and MICPCH_CASK_dup4/5_ (n=2). Active cells per cell line were pooled across replicates and statistical differences at 4 and 5 weeks were calculated using pairwise Wilcoxon Rank Sum test followed with Bonferroni correction.

### [1H]MRS

Proton magnetic resonance spectroscopy ([1H]MRS) data was available from the EU-AIMS Longitudinal European Autism Project (LEAP) autism twin cohort, including ASD_CASK_SS_ (Isaksson et al., 2018). MR data was acquired on MR750, 3 Tesla scanner (GE Healthcare, Milwaukie, USA), with 8 channel receiver array coil (in-VIVO inc.). To find small brain structures and to position the MRS voxel accurately and reproducibly a T1w 3D IR-SPGR (inversion-recovery spoiled gradient echo, resolution 1.0 x 1.0 x 1.2 mm) was acquired in the beginning of the protocol and the voxel position was confirmed thereafter by three-plane localizer images performed before every MRS scan. T1w MRI was setup according to ADNI2 protocol with flip angle 11°, te/tr = 3/ 7.3 ms, ti= 400 ms, number of slices 192 and bandwidth 244 Hz/px. MRS data was acquired with MEGA-PRESS pulse sequence, te/tr = 68/ 2000 ms, number of averages 192, phase cycle 8 steps and bandwidth 5 kHz for three voxels with volumes 19,3 mL (Medial PFC), 19.5 mL (Putamen-GPS), 13.6 mL (DLPFC) (Figure S6A). All voxels were positioned to maximize the grey matter (GM) content. CHESS water suppression and voxel OVS (6 outer volume suppression rf pulses) were applied. The gradient-echo shimming converged to the water linewidth of 10 ± 2 Hz for MPFC and Putamen-GPS and 6.9 ± 1 Hz for DLPFC.

GABA concentration of all three voxels was quantified from the MEGA-PRESS difference spectra using the “Gannet” GABA-MRS analysis toolkit, version 3.0 (Edden et al., 2014). Fitting of the GABA peak is performed over a range of the spectrum between 2.79 and 3.55 ppm using a five-parameter Gaussian model and of the water peak using a Gaussian-Lorentzian function. Measuring GABA using MEGA-PRESS leads to co-editing of macro-molecules such as proteins that contribute to the edited GABA peak at 3.0 ppm. Metabolite values were scaled to water and expressed in institutional units (IU). Water-scaled GABA concentrations represent GABA as well as related macromolecules and are therefore referred to as GABA+ concentrations (Figure S6B). GABA concentrations were adjusted for the average proportion of partial GM and white matter (WM) volume in each voxel across all subjects. For tissue correction, the GABA concentration ratio α between GM and WM was set to 0.5 (Harris et al., 2015). All spectra were visually evaluated for the quality of GABA fit. For the final analysis, 45 spectra from the Putamen (20 co-twins to individuals with NDDs), 33 from the MFC (11 co-twins to individuals with NDDs) and 44 from the DLPFC (19 co-twins to individuals with NDDs) of the control subjects could be included.

## Acknowledgments

The authors are grateful to the affected individuals and their families for their participation in this research. We also acknowledge the current and previous members of the RATSS team (Kerstin Andersson, Elodie Cauvet, Christina Coco, Johanna Ingvarsson, Elzbieta Kostrzewa, Johan Isaksson, Anna Råde, Annelies V’ant Westeinde, Elin Vahlgren, Karl Lundin, Martin Hammar, Lina Poltrago, Katell Mevel and Steve Berggren, at the Center of Neurodevelopmental Disorders at Karolinska Institutet (KIND)). The authors acknowledge support from the National Genomics Infrastructure in Stockholm funded by Science for Life Laboratory, the Knut and Alice Wallenberg Foundation and the Swedish Research Council, the SNIC/Uppsala Multidisciplinary Center for Advanced Computational Science for assistance with massively parallel sequencing and access to the UPPMAX computational infrastructure, the Eukaryotic Single-Cell Genomics facility at the Science for Life Laboratory for assistance with SMART-Seq2 sequencing, and the iPS Core facility at Karolinska Institutet for assistance with the generation of iPSC and NES cells.

The project was supported by the Swedish Research Council (E.H., A.F., S.B., K.T.), Swedish Foundation for Strategic Research (A.F., K.T.), The Swedish Brain Foundation – Hjärnfonden (E.H., A.F., S.B., K.T.), the Harald and Greta Jeanssons Foundations (K.T.), Åke Wiberg Foundation (K.T.), Jerring Foundation (S.B.), the Swedish Order of Freemasons (S.B), Kempe-Carlgrenska Foundation (S.B), Sunnderdahls Handikappsfond (S.B.), Solstickan Foundation (C.W), the Pediatric Research Foundation at Astrid Lindgren Children’s Hospital (S.B.), joint support from Swedish Research Council, Vinnova, Formas and FORTE (S.B), the Innovative Medicines Initiative Joint Undertaking (grant agreement number 115300), which comprises financial contribution from the European Union’s Seventh Framework Programme (FP7 /2007– 2013) and in-kind contributions from companies belonging to the European Federation of Pharmaceutical Industries and Associations(EU-AIMS), Strategic Research Area Neuroscience Stratneuro (K.T.), the L’Oréal-UNESCO for Women in Science prize in Sweden with support from the Young Academy of Sweden (K.T.), The Swedish Foundation for International Cooperation in Research and Higher Education STINT (K.T.) China Scholarship Council (D.L.), Queen Silvia’s Jubilee Fund (C.W., D.L.) the Frederik and Ingrid Thurings Foundation (M.B.) and Barnavård Foundation (M.B., D.L., K.T.) and Board of Research at Karolinska Institutet (K.T.).

## Author Contributions

M.B. and K.T. designed the study. J.W., C.W., L.M., E.H., S.B. and B.-M.A. performed acquisition of the phenotypic data, collection of biological samples and provided administrative support. S.M., M-L.H., J.N. and R.S. generated and analyzed the MR and/or MRS data. M.K. and A.F. generated iPSCs and NES cells and provided expertise in cell culture. M.B., F.M, I.R. V.M, L.B. and K.T performed genetic and molecular biology experiments. M.B. and F.M. performed immunostaining and confocal microscopy. M.B. and J.R. performed time-lapse calcium imaging with support from E.H.. M.B., F.M. and K.T. performed data and statistical analyses. E.H., A.F., S.B. and K.T. provided supervision in different parts of the study. All authors were part of interpreting the data and results. M.B., F.M., S.M., J.N., D.L. C.W. and K.T. prepared tables and figures. M.B and K.T drafted the manuscript. All authors performed a critical revision of the manuscript and approved the final version. M.B., S.B., A.F. and K.T. obtained funding.

## Supplementary Tables

**Table S1:** Psychometric profile of ASD_CASK_SS_ case and co-twin.

|  | **ASD<sub>CASK_SS</sub> case** | **Co-Twin** |
| --- | --- | --- |
| Early development | Language: Normal<br>Motoric: Normal<br>Toilet training: day-time enuresis until age 6y<br>First noted abnormalities: 4-5y | Language: Normal<br>Motoric: Normal<br>Toilet training: Normal<br>First noted abnormalities: 9y |
| Somatic | 1y: Colic and regurgitations<br>Eczema<br>From 2 years: Sleeping disorder –Insomnia<br>5y: Slight hearing impairment - right ear<br>From 5y: Visual impairment (hyperopia); prescription for glasses | 1y: Colic<br>Allergies<br>From 5y: Visual impairment (hyperopia; prescription for glasses) |
| Psychiatric diagnoses (DSM-IV) from Child and Youth Psychiatry | ADHD, Predominantly Inattentive Type (314.00)<br>PDD NOS (299.80) | ADHD, Predominantly Inattentive Type (314.00)<br>PDD NOS (299.80) |
| Intellectual level | GAI: 121, VCI: 106, PCI: 129 | GAI: 121, VCI: 110, PCI: 125 |
| Adaptive functioning | GAF: 87 | GAF: 82 |
| ADHD deficits | Global Index: 7; AN: 7; AH: 7 | Global Index: 7; AN: 11; AH: 4 |
| Autism symptoms | AQ: 73<br>SRS total score: 69<br>ADI-R: 11-7-3-0<br>ADOS total: 2-4-1-0 | AQ: 60<br>SRS total score: 53<br>ADI-R: 11-7-0-0<br>ADOS total: 2-3-2-0 |
| Executive functioning | The Tower total scale score: 10<br>WCST total errors t-score: 61 | The Tower total scale score: 11<br>WCST total errors t-score: 65 |
| Psychiatric symptoms | Insomnia, Tics, Anxiety | - |
| Medications at time for assessment | Concerta, Strattera | No medication |
| Morphological features | Hypermobility in fingers<br>Long eyelashes<br>Hypopigmentation (i.e., vitiligo)<br>Inverted mamillary glands<br>Abnormal tear production- tears easily<br>Vision impairment<br>Overbite (teeth)<br>Sloping shoulders<br>Light hair | Overweight<br>Hitchhiker thumb<br>Long eyelashes<br>Vision impairment<br>Hypertelorism<br>Long palpebral fissure<br>Abnormal tear production- tears easily<br>High palate<br>Overbite (teeth)<br>Small ears<br>Preauricular skin tag<br>Light hair<br>Sandal gap |

## Supplementary Figures

**Figure S1.**
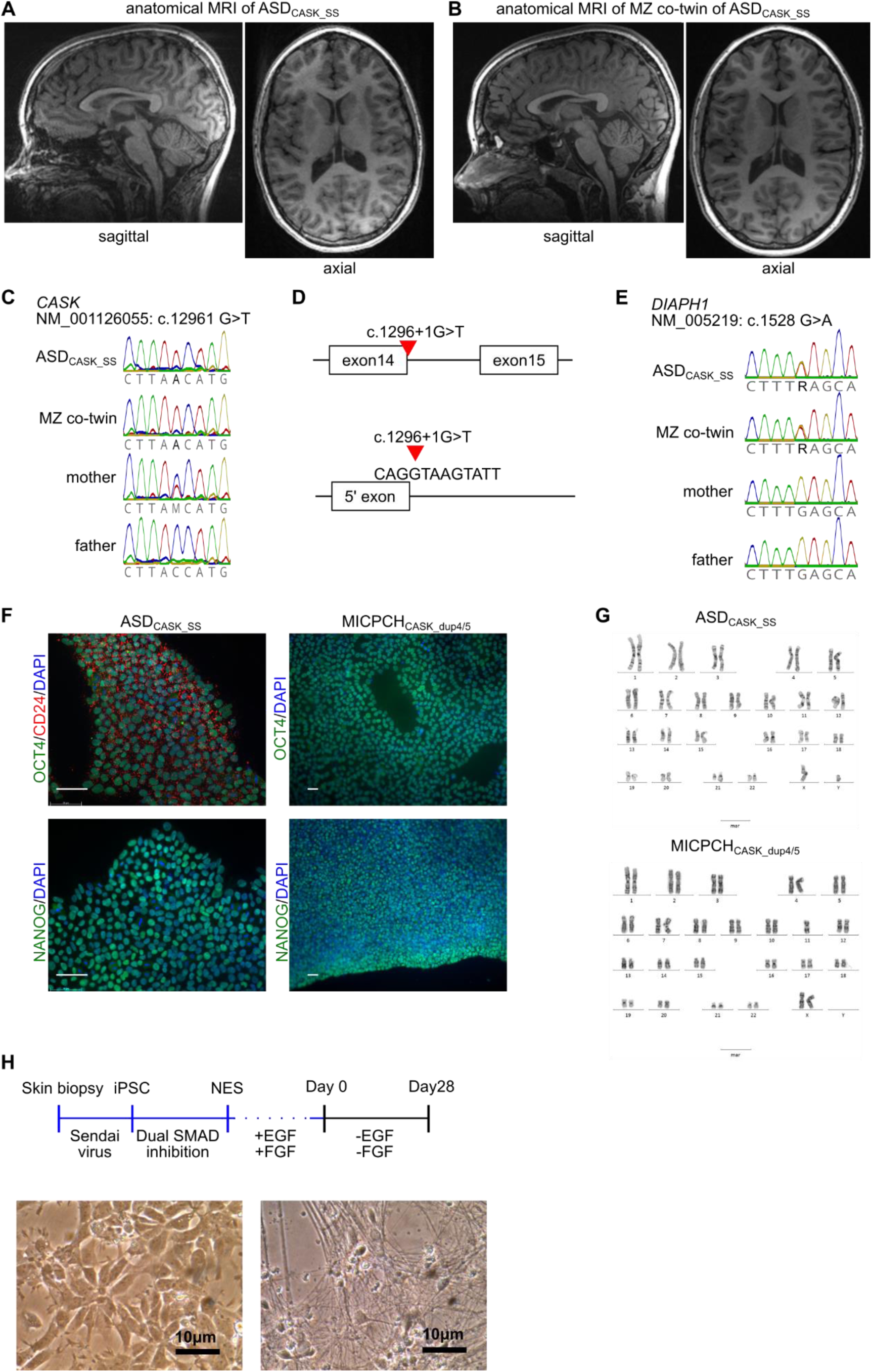
(A, B) Sagittal and axial anatomical magnetic resonance images of (A) ASD_CASK_SS_ and (B) his monozygotic co-twin. (C) Sanger sequencing validation of the CASK variant NM_001126055: c.1296+1 G>T detected in whole exome sequencing. Sequencing was performed on genomic DNA obtained from blood of individual ASD_CASK_SS_, his co-twin and parents. (D) Schematic location of the CASK splice site variant between exon 14 and intron 14. The mutation changes the +1 guanine (G) to a thymine (T), which is not represented in the position weight matrix of donor splice sites. (E) Sanger sequencing validation of the DIAPH1 variant NM_005219: c.1528 G>A detected in whole exome sequencing. Sequencing was performed on genomic DNA obtained from blood of individual ASD_CASK_SS_, his co-twin and parents. (F) Pluripotency marker NANOG and OCT4 staining in iPSC cells derived from individuals ASD_CASK_SS_ and MICPCH_CASK_dup4/5_. Scale = 50 µm. (G) Karyotypes of iPSC clones obtained from ASD_CASK_SS_ and MICPCH_CASK_dup_4/5_. (H) Schematic summary of differentiation protocol from fibroblasts to maturing neurons and representative phase-contrast microscopy images of male control NES cells and day 28 neurons.

**Figure S2.**
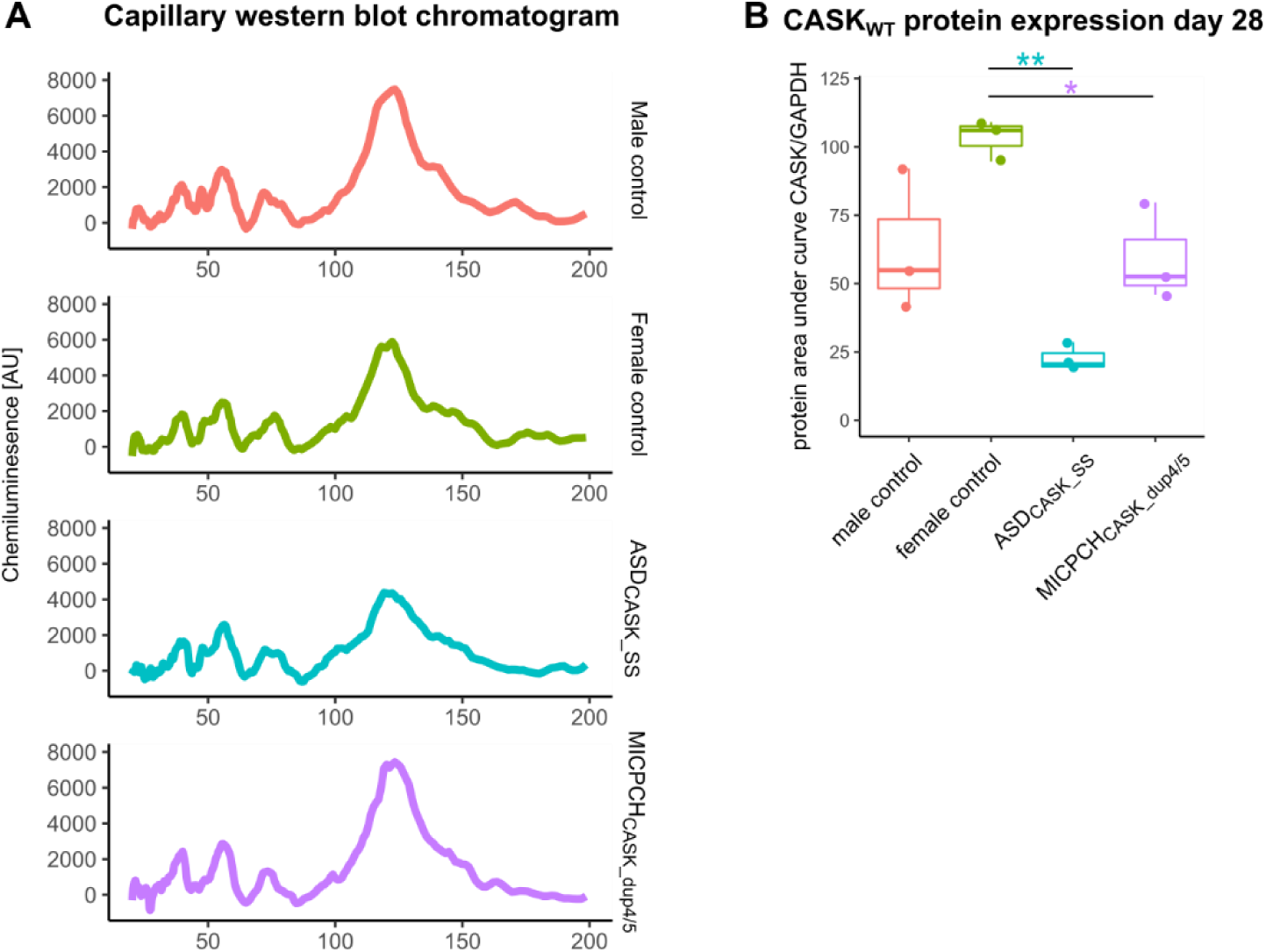
(A) Chromatogram of protein samples stained with CASK antibody and run on capillary western blot. (B) CASK mRNA and protein expression at 28 days of neuronal differentiation. Statistical difference was calculated with ANOVA and post-hoc Tukey. *<0.05, **<0.01, ***<0.001.

**Figure S3.**
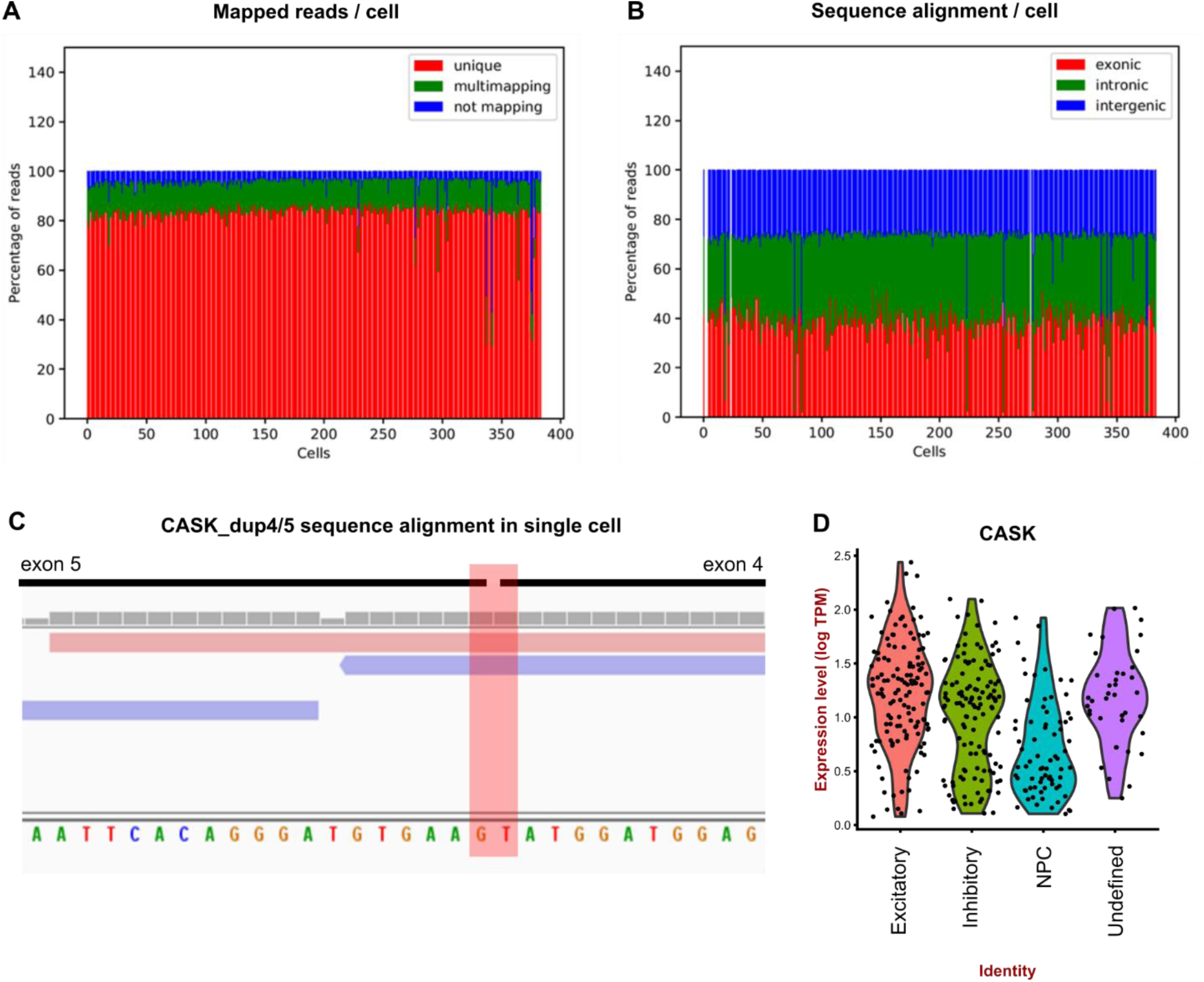
(A) Detection of CASK_dup4_5_ variant in SMART-Seq2 sequencing of one cell shown in IGV browser. In this cell, two reads align to the unique exon5 to exon 4 junction of the MICPCH_CASK_dup4/5_ case. (B) Violin plot of CASK expression in all sequenced cells grouped into identified cell-types.

**Figure S4.**
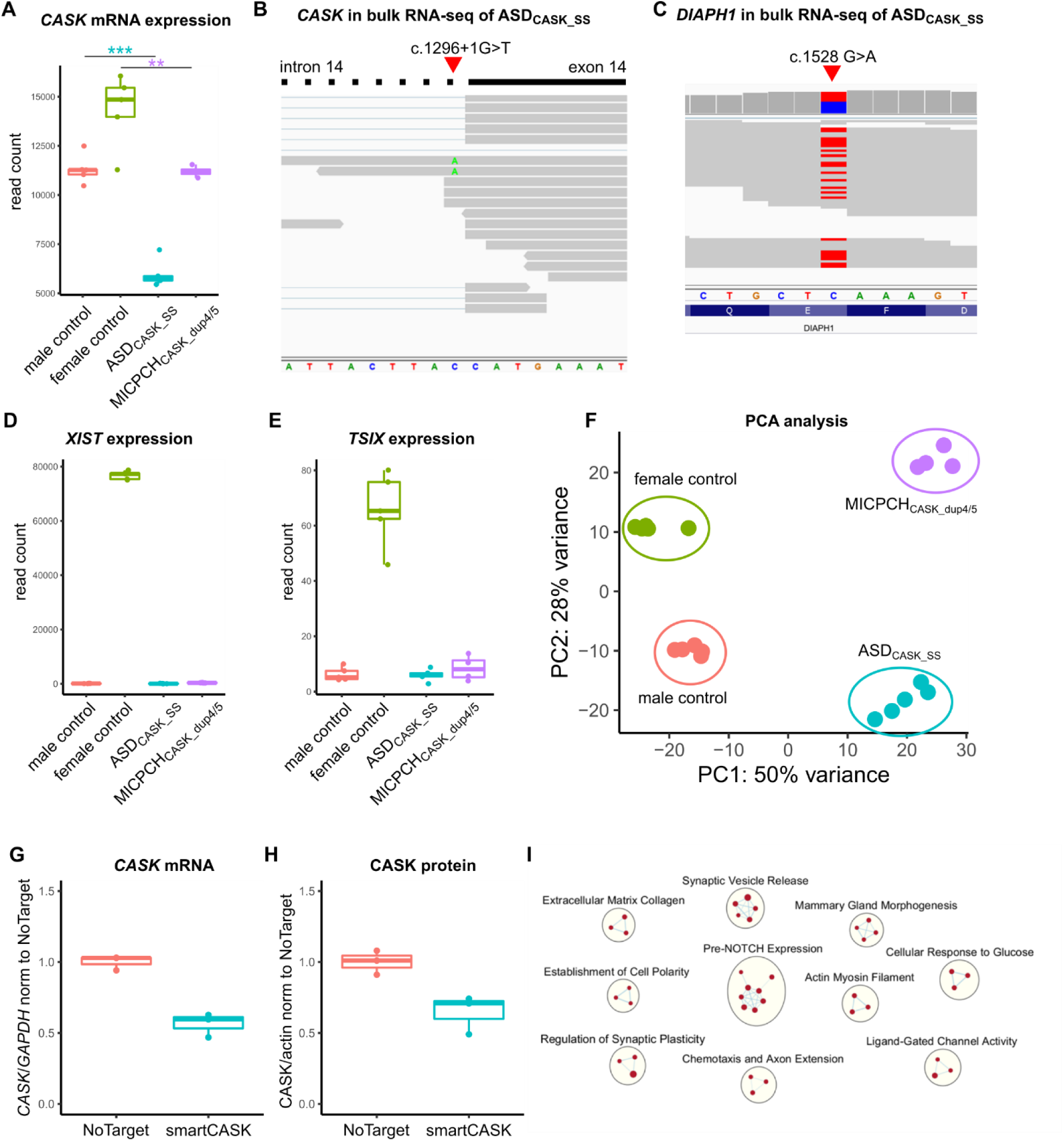
(A) CASK mRNA expression in bulk RNA samples of cases and controls validates qPCR expression results. (B) Detection of RNA Sequence reads at the exon14-intron14 boundary of one ASD_CASK_SS_ sample. Sequence reads provide evidence for CASK_WT_ and CASK_14+_ with reference G and mutated T nucleotide. (C) Detection of *de novo* DIAPH1 variant in RNA sequence reads of ASD_CASK_SS_ showing biallelic expression of wildtype and mutant allele. (D, E) Expression of *XIST* and *TSIX* non-coding RNAs in cases and controls showing high expression only in the female control cell line. (F) PCA of bulk RNA-sequence samples indicating separation of cases and control along PC1 and males and females along PC2. (G, H) Expression of CASK mRNA and protein after siRNA mediated knockdown indicating 60-70% remaining CASK expression in comparison to a no-target siRNA control. (I) GSEA enriched pathways of upregulated genes after siRNA knockdown.

**Figure S5.**
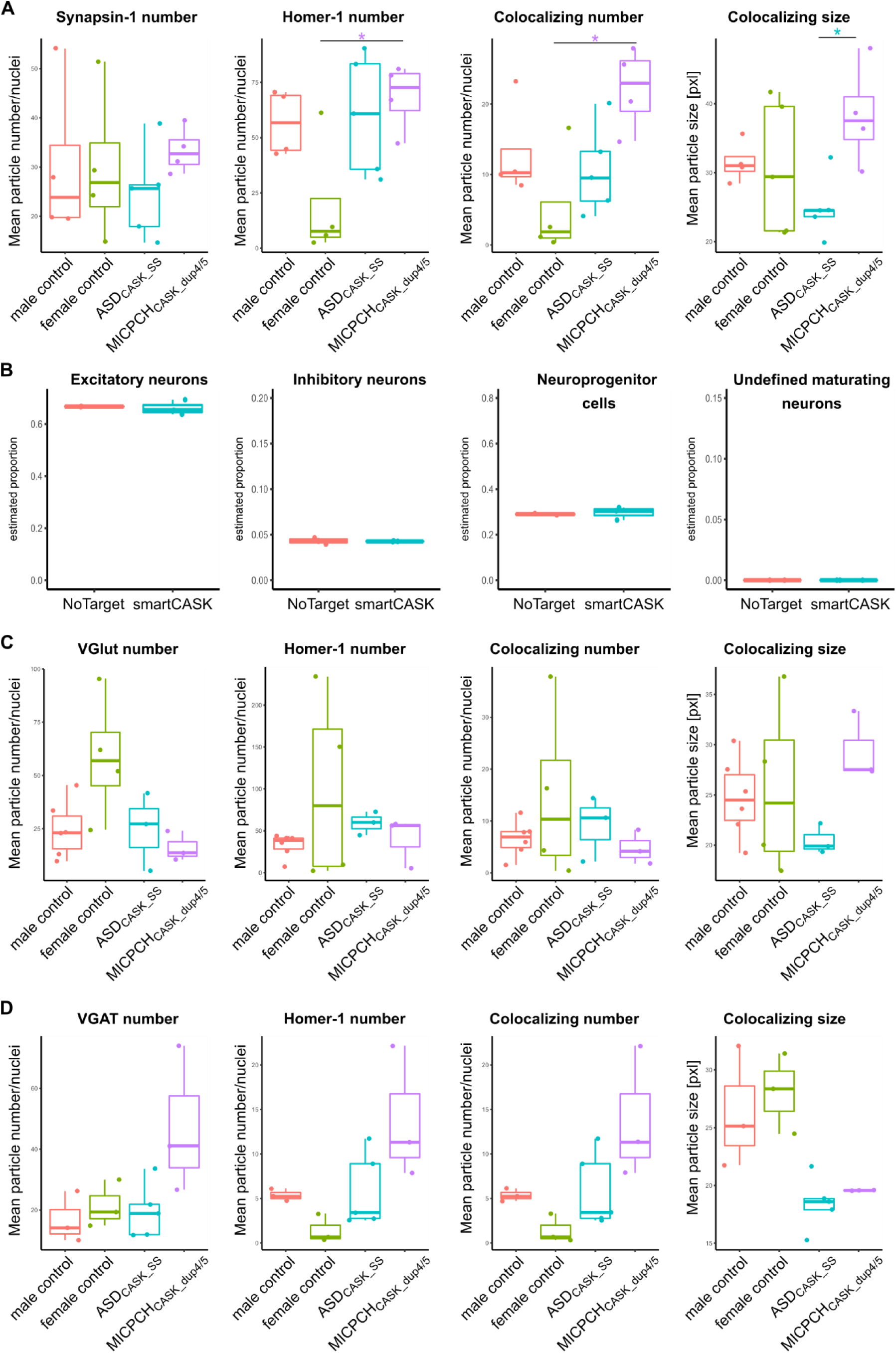
(A) Synapse Counter results for Synapsin-1/Homer-1 co-staining of case and control cell lines differentiated for 28 days. The panel of graphs shows the Synapsin-1 and Homer-1 number per nuclei, the number of co-localizing region and the mean size of co-localizing sizes per biological sample. Number of replicates and statistical analysis as stated in legend of Figure 5. (B) Deconvolution of cell populations in siRNA knockdown RNA sequencing samples from MICPCH_CASK_dup4/5_ single-cell RNA-sequencing dataset. (C) Synapse Counter results for VGlut/Homer-1 co-staining of case and control cell lines differentiated for 28 days. The panel of graphs shows the VGlut and Homer-1 number per nuclei, the number of co-localizing region and the mean size of co-localizing sizes per biological sample. Number of replicates and statistical analysis as stated in legend of Figure 6. (D) Synapse Counter results for VGAT/Homer-1 co-staining of case and control cell lines differentiated for 28 days. The panel of graphs shows the VGAT and Homer-1 number per nuclei, the number of co-localizing region and the mean size of co-localizing sizes per biological sample. Number of replicates and statistical analysis as stated in legend of Figure 6.

**Figure S6.**
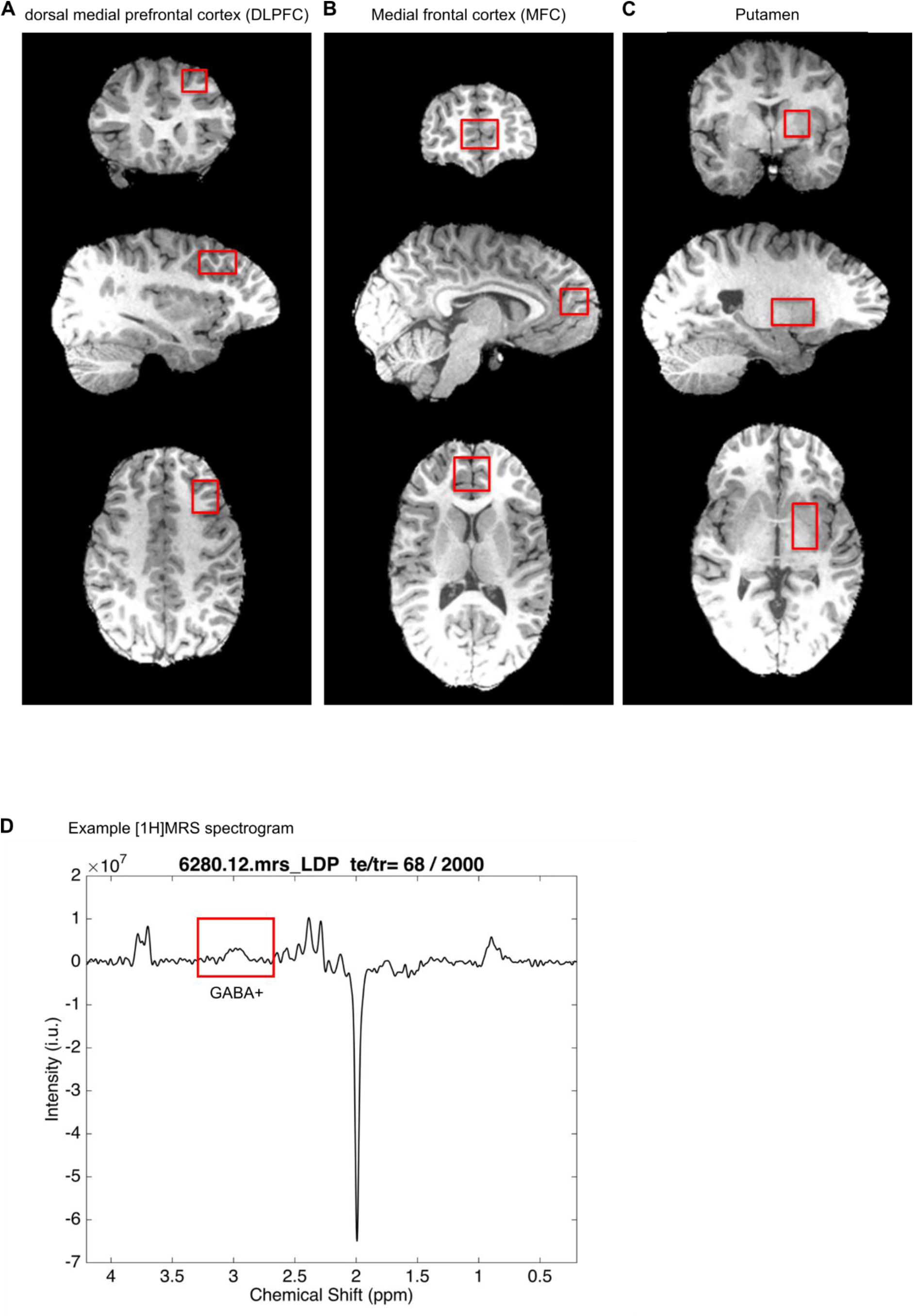
(A-C) Example MR images underlying [1H]MRS, indicating the voxel regions used to detect GABA concentrations in (A) the DLPFC, (B) MFC and (C) putamen. (D) Example [1H]MRS spectrum obtained from a 13.6 ml voxel located at the DLPFC, indicating the spectral peak of GABA.

## References

Al-Maawali, A., Barry, B.J., Rajab, A., El-Quessny, M., Seman, A., Coury, S.N., Barkovich, A.J., Yang, E., Walsh, C.A., Mochida, G.H., et al. (2016). Novel loss-of-function variants in DIAPH1 associated with syndromic microcephaly, blindness, and early onset seizures. American journal of medical genetics Part A 170A, 435–440.

Al Shehhi, M., Forman, E.B., Fitzgerald, J.E., McInerney, V., Krawczyk, J., Shen, S., Betts, D.R., Ardle, L.M., Gorman, K.M., King, M.D., et al. (2019). NRXN1 deletion syndrome; phenotypic and penetrance data from 34 families. European journal of medical genetics 62, 204–209.

Amatya, D.N., Linker, S.B., Mendes, A.P.D., Santos, R., Erikson, G., Shokhirev, M.N., Zhou, Y.S., Sharpee, T., Gage, F.H., Marchetto, M.C., et al. (2019). Dynamical Electrical Complexity Is Reduced during Neuronal Differentiation in Autism Spectrum Disorder. Stem cell reports 13, 474–484.

Arora, M., Reichenberg, A., Willfors, C., Austin, C., Gennings, C., Berggren, S., Lichtenstein, P., Anckarsater, H., Tammimies, K., and Bolte, S. (2017). Fetal and postnatal metal dysregulation in autism. Nature communications 8, 15493.

Artimovich, E., Jackson, R.K., Kilander, M.B.C., Lin, Y.C., and Nestor, M.W. (2017). PeakCaller: an automated graphical interface for the quantification of intracellular calcium obtained by high-content screening. Bmc Neurosci 18.

Atasoy, D., Schoch, S., Ho, A., Nadasy, K.A., Liu, X.R., Zhang, W.Q., Mukherjee, K., Nosyreva, E.D., Fernandez-Chacon, R., Missler, M., et al. (2007). Deletion of CASK in mice is lethal and impairs synaptic function. Proc Natl Acad Sci U S A 104, 2525–2530.

Baker, K., Gordon, S.L., Melland, H., Bumbak, F., Scott, D.J., Jiang, T.J., Owen, D., Turner, B.J., Boyd, S.G., Rossi, M., et al. (2018). SYT1-associated neurodevelopmental disorder: a case series. Brain : a journal of neurology 141, 2576–2591.

Bakkaloglu, B., O’Roak, B.J., Louvi, A., Gupta, A.R., Abelson, J.F., Morgan, T.M., Chawarska, K., Klin, A., Ercan-Sencicek, A.G., Stillman, A.A., et al. (2008). Molecular cytogenetic analysis and resequencing of contactin associated protein-like 2 in autism spectrum disorders. Am J Hum Genet 82, 165–173.

Baron, M., Veres, A., Wolock, S.L., Faust, A.L., Gaujoux, R., Vetere, A., Ryu, J.H., Wagner, B.K., Shen-Orr, S.S., Klein, A.M., et al. (2016). A Single-Cell Transcriptomic Map of the Human and Mouse Pancreas Reveals Inter- and Intra-cell Population Structure. Cell Syst 3, 346–360 e344.

Ben-Ari, Y. (2002). Excitatory actions of gaba during development: the nature of the nurture. Nat Rev Neurosci 3, 728–739.

Bölte, S., Willfors, C., Berggren, S., Norberg, J., Poltrago, L., Mevel, K., Coco, C., Fransson, P., Borg, J., Sitnikov, R., et al. (2014). The Roots of Autism and ADHD Twin Study in Sweden (RATSS). Twin Res Hum Genet 17, 164–176.

Burglen, L., Chantot-Bastaraud, S., Garel, C., Milh, M., Touraine, R., Zanni, G., Petit, F., Afenjar, A., Goizet, C., Barresi, S., et al. (2012). Spectrum of pontocerebellar hypoplasia in 13 girls and boys with CASK mutations: confirmation of a recognizable phenotype and first description of a male mosaic patient. Orphanet J Rare Dis 7, 18.

Butler, A., Hoffman, P., Smibert, P., Papalexi, E., and Satija, R. (2018). Integrating single-cell transcriptomic data across different conditions, technologies, and species. Nat Biotechnol 36, 411–420.

Butz, S., Okamoto, M., and Sudhof, T.C. (1998). A tripartite protein complex with the potential to couple synaptic vesicle exocytosis to cell adhesion in brain. Cell 94, 773–782.

Carithers, L.J., Ardlie, K., Barcus, M., Branton, P.A., Britton, A., Buia, S.A., Compton, C.C., DeLuca, D.S., Peter-Demchok, J., Gelfand, E.T., et al. (2015). A Novel Approach to High-Quality Postmortem Tissue Procurement: The GTEx Project. Biopreserv Biobank 13, 311–319.

Chambers, S.M., Fasano, C.A., Papapetrou, E.P., Tomishima, M., Sadelain, M., and Studer, L. (2009). Highly efficient neural conversion of human ES and iPS cells by dual inhibition of SMAD signaling. Nat Biotechnol 27, 275–280.

Coe, B.P., Stessman, H.A.F., Sulovari, A., Geisheker, M.R., Bakken, T.E., Lake, A.M., Dougherty, J.D., Lein, E.S., Hormozdiari, F., Bernier, R.A., et al. (2019). Neurodevelopmental disease genes implicated by de novo mutation and copy number variation morbidity. Nat Genet 51, 106–116.

Cristofoli, F., Devriendt, K., Davis, E.E., Van Esch, H., and Vermeesch, J.R. (2018). Novel CASK mutations in cases with syndromic microcephaly. Human mutation 39, 993–1001.

Dabell, M.P., Rosenfeld, J.A., Bader, P., Escobar, L.F., El-Khechen, D., Vallee, S.E., Dinulos, M.B.P., Curry, C., Fisher, J., Tervo, R., et al. (2013). Investigation of NRXN1 deletions: Clinical and molecular characterization. American Journal of Medical Genetics Part A 161a, 717–731.

Dai, D.P., Gan, W., Hayakawa, H., Zhu, J.L., Zhang, X.Q., Hu, G.X., Xu, T., Jiang, Z.L., Zhang, L.Q., Hu, X.D., et al. (2018). Transcriptional mutagenesis mediated by 8-oxoG induces translational errors in mammalian cells. Proc Natl Acad Sci U S A 115, 4218–4222.

Dandulakis, M.G., Meganathan, K., Kroll, K.L., Bonni, A., and Constantino, J.N. (2016). Complexities of X chromosome inactivation status in female human induced pluripotent stem cells-a brief review and scientific update for autism research. J Neurodev Disord 8.

De Rubeis, S., He, X., Goldberg, A.P., Poultney, C.S., Samocha, K., Cicek, A.E., Kou, Y., Liu, L., Fromer, M., Walker, S., et al. (2014). Synaptic, transcriptional and chromatin genes disrupted in autism. Nature 515, 209–U119.

Deciphering Developmental Disorders, S. (2017). Prevalence and architecture of de novo mutations in developmental disorders. Nature 542, 433–438.

DeLuca, S.C., Wallace, D.A., Trucks, M.R., and Mukherjee, K. (2017). A clinical series using intensive neurorehabilitation to promote functional motor and cognitive skills in three girls with CASK mutation. BMC Research Notes 10, 743.

Deneault, E., White, S.H., Rodrigues, D.C., Ross, P.J., Faheem, M., Zaslavsky, K., Wang, Z., Alexandrova, R., Pellecchia, G., Wei, W., et al. (2018). Complete Disruption of Autism-Susceptibility Genes by Gene Editing Predominantly Reduces Functional Connectivity of Isogenic Human Neurons. Stem cell reports 11, 1211–1225.

Deriziotis, P., O’Roak, B.J., Graham, S.A., Estruch, S.B., Dimitropoulou, D., Bernier, R.A., Gerdts, J., Shendure, J., Eichler, E.E., and Fisher, S.E. (2014). De novo TBR1 mutations in sporadic autism disrupt protein functions. Nature communications 5, 4954.

Dias, C., Estruch, S.B., Graham, S.A., McRae, J., Sawiak, S.J., Hurst, J.A., Joss, S.K., Holder, S.E., Morton, J.E., Turner, C., et al. (2016). BCL11A Haploinsufficiency Causes an Intellectual Disability Syndrome and Dysregulates Transcription. Am J Hum Genet 99, 253–274.

Dick, A.L.W., Khermesh, K., Paul, E., Stamp, F., Levanon, E.Y., and Chen, A. (2019). Adenosine-to-Inosine RNA Editing Within Corticolimbic Brain Regions Is Regulated in Response to Chronic Social Defeat Stress in Mice. Frontiers in Psychiatry 10.

Dzyubenko, E., Rozenberg, A., Hermann, D.M., and Faissner, A. (2016). Colocalization of synapse marker proteins evaluated by STED-microscopy reveals patterns of neuronal synapse distribution in vitro. J Neurosci Methods 273, 149–159.

Edden, R.A.E., Puts, N.A.J., Harris, A.D., Barker, P.B., and Evans, C.J. (2014). Gannet: A Batch-Processing Tool for the Quantitative Analysis of Gamma-Aminobutyric Acid-Edited MR Spectroscopy Spectra. J Magn Reson Imaging 40, 1445–1452.

Edelstein, A.D., Tsuchida, M.A., Amodaj, N., Pinkard, H., Vale, R.D., and Stuurman, N. (2014). Advanced methods of microscope control using muManager software. J Biol Methods 1.

Ercan-Sencicek, A.G., Jambi, S., Franjic, D., Nishimura, S., Li, M., El-Fishawy, P., Morgan, T.M., Sanders, S.J., Bilguvar, K., Suri, M., et al. (2015). Homozygous loss of DIAPH1 is a novel cause of microcephaly in humans. European journal of human genetics : EJHG 23, 165–172.

Erickson, C.A., Davenport, M.H., Schaefer, T.L., Wink, L.K., Pedapati, E.V., Sweeney, J.A., Fitzpatrick, S.E., Brown, W.T., Budimirovic, D., Hagerman, R.J., et al. (2017). Fragile X targeted pharmacotherapy: lessons learned and future directions. J Neurodev Disord 9.

Erickson, C.A., Veenstra-Vanderweele, J.M., Melmed, R.D., McCracken, J.T., Ginsberg, L.D., Sikich, L., Scahill, L., Cherubini, M., Zarevics, P., Walton-Bowen, K., et al. (2014a). STX209 (arbaclofen) for autism spectrum disorders: an 8-week open-label study. J Autism Dev Disord 44, 958–964.

Erickson, C.A., Wink, L.K., Early, M.C., Stiegelmeyer, E., Mathieu-Frasier, L., Patrick, V., and McDougle, C.J. (2014b). Brief report: Pilot single-blind placebo lead-in study of acamprosate in youth with autistic disorder. J Autism Dev Disord 44, 981–987.

Etherton, M.R., Blaiss, C.A., Powell, C.M., and Sudhof, T.C. (2009). Mouse neurexin-1alpha deletion causes correlated electrophysiological and behavioral changes consistent with cognitive impairments. Proc Natl Acad Sci U S A 106, 17998–18003.

Falk, A., Koch, P., Kesavan, J., Takashima, Y., Ladewig, J., Alexander, M., Wiskow, O., Tailor, J., Trotter, M., Pollard, S., et al. (2012). Capture of neuroepithelial-like stem cells from pluripotent stem cells provides a versatile system for in vitro production of human neurons. PloS one 7, e29597.

Ford, T.C., and Crewther, D.P. (2016). A Comprehensive Review of the (1)H-MRS Metabolite Spectrum in Autism Spectrum Disorder. Front Mol Neurosci 9, 14.

Forsberg, D., Thonabulsombat, C., Jaderstad, J., Jaderstad, L.M., Olivius, P., and Herlenius, E. (2017). Functional Stem Cell Integration into Neural Networks Assessed by Organotypic Slice Cultures. Curr Protoc Stem Cell Biol 42, 2D 13 11-12D 13 30.

Gao, R., Piguel, N.H., Melendez-Zaidi, A.E., Martin-de-Saavedra, M.D., Yoon, S., Forrest, M.P., Myczek, K., Zhang, G., Russell, T.A., Csernansky, J.G., et al. (2018). CNTNAP2 stabilizes interneuron dendritic arbors through CASK. Molecular psychiatry 23, 1832–1850.

Gao, R., Zaccard, C.R., Shapiro, L.P., Dionisio, L.E., Martin-de-Saavedra, M.D., Piguel, N.H., Pratt, C.P., Horan, K.E., and Penzes, P. (2019). The CNTNAP2-CASK complex modulates GluA1 subcellular distribution in interneurons. Neuroscience letters 701, 92–99.

Gupta, A.R., Pirruccello, M., Cheng, F., Kang, H.J., Fernandez, T.V., Baskin, J.M., Choi, M., Liu, L., Ercan-Sencicek, A.G., Murdoch, J.D., et al. (2014). Rare deleterious mutations of the gene EFR3A in autism spectrum disorders. Molecular autism 5, 31.

Hackett, A., Tarpey, P.S., Licata, A., Cox, J., Whibley, A., Boyle, J., Rogers, C., Grigg, J., Partington, M., Stevenson, R.E., et al. (2010). CASK mutations are frequent in males and cause X-linked nystagmus and variable XLMR phenotypes. European Journal of Human Genetics 18, 544–552.

Harris, A.D., Puts, N.A.J., and Edden, R.A.E. (2015). Tissue correction for GABA-edited MRS: Considerations of voxel composition, tissue segmentation, and tissue relaxations. J Magn Reson Imaging 42, 1431–1440.

Hata, Y., Butz, S., and Sudhof, T.C. (1996). CASK: a novel dlg/PSD95 homolog with an N-terminal calmodulin-dependent protein kinase domain identified by interaction with neurexins. The Journal of neuroscience : the official journal of the Society for Neuroscience 16, 2488–2494.

Hayashi, S., Uehara, D.T., Tanimoto, K., Mizuno, S., Chinen, Y., Fukumura, S., Takanashi, J.I., Osaka, H., Okamoto, N., and Inazawa, J. (2017). Comprehensive investigation of CASK mutations and other genetic etiologies in 41 patients with intellectual disability and microcephaly with pontine and cerebellar hypoplasia (MICPCH). PloS one 12, e0181791.

Hsueh, Y.P., and Sheng, M. (1999). Regulated expression and subcellular localization of syndecan heparan sulfate proteoglycans and the syndecan-binding protein CASK/LIN-2 during rat brain development. The Journal of neuroscience : the official journal of the Society for Neuroscience 19, 7415–7425.

Hu, H.T., Umemori, H., and Hsueh, Y.P. (2016). Postsynaptic SDC2 induces transsynaptic signaling via FGF22 for bidirectional synaptic formation. Sci Rep 6.

Iossifov, I., O’Roak, B.J., Sanders, S.J., Ronemus, M., Krumm, N., Levy, D., Stessman, H.A., Witherspoon, K.T., Vives, L., Patterson, K.E., et al. (2014). The contribution of de novo coding mutations to autism spectrum disorder. Nature 515, 216–U136.

Isaksson, J., Tammimies, K., Neufeld, J., Cauvet, E., Lundin, K., Buitelaar, J.K., Loth, E., Murphy, D.G.M., Spooren, W., Bolte, S., et al. (2018). EU-AIMS Longitudinal European Autism Project (LEAP): the autism twin cohort. Molecular autism 9, 26.

Jia, H., Rochefort, N.L., Chen, X., and Konnerth, A. (2011). In vivo two-photon imaging of sensory-evoked dendritic calcium signals in cortical neurons. Nature protocols 6, 28–35.

Karczewski, K.J., Francioli, L.C., Tiao, G., Cummings, B.B., Alföldi, J., Wang, Q., Collins, R.L., Laricchia, K.M., Ganna, A., Birnbaum, D.P., et al. (2019). Variation across 141,456 human exomes and genomes reveals the spectrum of loss-of-function intolerance across human protein-coding genes. bioRxiv.

Kohler, S., Doelken, S.C., Mungall, C.J., Bauer, S., Firth, H.V., Bailleul-Forestier, I., Black, G.C.M., Brown, D.L., Brudno, M., Campbell, J., et al. (2014). The Human Phenotype Ontology project: linking molecular biology and disease through phenotype data. Nucleic Acids Res 42, D966–D974.

Kuo, T.Y., Hong, C.J., Chien, H.L., and Hsueh, Y.P. (2010). X-linked mental retardation gene CASK interacts with Bcl11A/CTIP1 and regulates axon branching and outgrowth. Journal of neuroscience research 88, 2364–2373.

LaConte, L.E., Chavan, V., Liang, C., Willis, J., Schonhense, E.M., Schoch, S., and Mukherjee, K. (2016). CASK stabilizes neurexin and links it to liprin-alpha in a neuronal activity-dependent manner. Cell Mol Life Sci 73, 3599–3621.

LaConte, L.E., Chavan, V., and Mukherjee, K. (2014). Identification and glycerol-induced correction of misfolding mutations in the X-linked mental retardation gene CASK. PloS one 9, e88276.

LaConte, L.E.W., Chavan, V., Elias, A.F., Hudson, C., Schwanke, C., Styren, K., Shoof, J., Kok, F., Srivastava, S., and Mukherjee, K. (2018). Two microcephaly-associated novel missense mutations in CASK specifically disrupt the CASK-neurexin interaction. Human genetics 137, 231–246.

Lam, M., Moslem, M., Bryois, J., Pronk, R.J., Uhlin, E., Ellstrom, I.D., Laan, L., Olive, J., Morse, R., Ronnholm, H., et al. (2019). Single cell analysis of autism patient with bi-allelic NRXN1-alpha deletion reveals skewed fate choice in neural progenitors and impaired neuronal functionality. Exp Cell Res 383.

Li, W.V., and Li, J.J. (2018). An accurate and robust imputation method scImpute for single-cell RNA-seq data. Nature communications 9, 997.

Li, Y.C., and Kavalali, E.T. (2017). Synaptic Vesicle-Recycling Machinery Components as Potential Therapeutic Targets. Pharmacol Rev 69, 141–160.

Lin, E.I., Jeyifous, O., and Green, W.N. (2013). CASK regulates SAP97 conformation and its interactions with AMPA and NMDA receptors. The Journal of neuroscience : the official journal of the Society for Neuroscience 33, 12067–12076.

Lynch, E.D., Lee, M.K., Morrow, J.E., Welcsh, P.L., Leon, P.E., and King, M.C. (1997). Nonsyndromic deafness DFNA1 associated with mutation of a human homolog of the Drosophila gene diaphanous. Science 278, 1315–1318.

Marchetto, M.C., Belinson, H., Tian, Y., Freitas, B.C., Fu, C., Vadodaria, K.C., Beltrao-Braga, P.C., Trujillo, C.A., Mendes, A.P.D., Padmanabhan, K., et al. (2017). Altered proliferation and networks in neural cells derived from idiopathic autistic individuals. Molecular psychiatry 22, 820–835.

Merico, D., Isserlin, R., Stueker, O., Emili, A., and Bader, G.D. (2010). Enrichment Map: A Network-Based Method for Gene-Set Enrichment Visualization and Interpretation. PloS one 5.

Moey, C., Hinze, S.J., Brueton, L., Morton, J., McMullan, D.J., Kamien, B., Barnett, C.P., Brunetti-Pierri, N., Nicholl, J., Gecz, J., et al. (2016). Xp11.2 microduplications including IQSEC2, TSPYL2 and KDM5C genes in patients with neurodevelopmental disorders. European journal of human genetics : EJHG 24, 373–380.

Moog, U., Bierhals, T., Brand, K., Bautsch, J., Biskup, S., Brune, T., Denecke, J., de Die-Smulders, C.E., Evers, C., Hempel, M., et al. (2015). Phenotypic and molecular insights into CASK-related disorders in males. Orphanet J Rare Dis 10.

Moog, U., Kutsche, K., Kortum, F., Chilian, B., Bierhals, T., Apeshiotis, N., Balg, S., Chassaing, N., Coubes, C., Das, S., et al. (2011). Phenotypic spectrum associated with CASK loss-of-function mutations. Journal of medical genetics 48, 741–751.

Mori, T., Kasem, E.A., Suzuki-Kouyama, E., Cao, X., Li, X., Kurihara, T., Uemura, T., Yanagawa, T., and Tabuchi, K. (2019). Deficiency of calcium/calmodulin-dependent serine protein kinase disrupts the excitatory-inhibitory balance of synapses by down-regulating GluN2B. Molecular psychiatry.

Muller, F.J., Schuldt, B.M., Williams, R., Mason, D., Altun, G., Papapetrou, E.P., Danner, S., Goldmann, J.E., Herbst, A., Schmidt, N.O., et al. (2011). A bioinformatic assay for pluripotency in human cells. Nat Methods 8, 315–U354.

Myers, L., Anderlid, B.M., Nordgren, A., Willfors, C., Kuja-Halkola, R., Tammimies, K., and Bolte, S. (2017). Minor physical anomalies in neurodevelopmental disorders: a twin study. Child Adolesc Psychiatry Ment Health 11, 57.

Najm, J., Horn, D., Wimplinger, I., Golden, J.A., Chizhikov, V.V., Sudi, J., Christian, S.L., Ullmann, R., Kuechler, A., Haas, C.A., et al. (2008). Mutations of CASK cause an X-linked brain malformation phenotype with microcephaly and hypoplasia of the brainstem and cerebellum. Nat Genet 40, 1065–1067.

Neuhaus, C., Lang-Roth, R., Zimmermann, U., Heller, R., Eisenberger, T., Weikert, M., Markus, S., Knipper, M., and Bolz, H.J. (2017). Extension of the clinical and molecular phenotype of DIAPH1-associated autosomal dominant hearing loss (DFNA1). Clin Genet 91, 892–901.

O’Roak, B.J., Deriziotis, P., Lee, C., Vives, L., Schwartz, J.J., Girirajan, S., Karakoc, E., Mackenzie, A.P., Ng, S.B., Baker, C., et al. (2011). Exome sequencing in sporadic autism spectrum disorders identifies severe de novo mutations. Nature genetics 43, 585–589.

Pachitariu, M., Stringer, C., and Harris, K.D. (2018). Robustness of Spike Deconvolution for Neuronal Calcium Imaging. The Journal of neuroscience : the official journal of the Society for Neuroscience 38, 7976–7985.

Pak, C., Danko, T., Zhang, Y., Aoto, J., Anderson, G., Maxeiner, S., Yi, F., Wernig, M., and Sudhof, T.C. (2015). Human Neuropsychiatric Disease Modeling using Conditional Deletion Reveals Synaptic Transmission Defects Caused by Heterozygous Mutations in NRXN1. Cell stem cell 17, 316–328.

Picelli, S., Bjorklund, A.K., Faridani, O.R., Sagasser, S., Winberg, G., and Sandberg, R. (2013). Smart-seq2 for sensitive full-length transcriptome profiling in single cells. Nat Methods 10, 1096–1098.

Piluso, G., D’Amico, F., Saccone, V., Bismuto, E., Rotundo, I.L., Di Domenico, M., Aurino, S., Schwartz, C.E., Neri, G., and Nigro, V. (2009). A missense mutation in CASK causes FG syndrome in an Italian family. Am J Hum Genet 84, 162–177.

Pinto, D., Delaby, E., Merico, D., Barbosa, M., Merikangas, A., Klei, L., Thiruvahindrapuram, B., Xu, X., Ziman, R., Wang, Z., et al. (2014). Convergence of genes and cellular pathways dysregulated in autism spectrum disorders. Am J Hum Genet 94, 677–694.

Plummer, J.T., Gordon, A.J., and Levitt, P. (2016). The Genetic Intersection of Neurodevelopmental Disorders and Shared Medical Comorbidities - Relations that Translate from Bench to Bedside. Front Psychiatry 7, 142.

Reid, J.G., Nagaraja, A.K., Lynn, F.C., Drabek, R.B., Muzny, D.M., Shaw, C.A., Weiss, M.K., Naghavi, A.O., Khan, M., Zhu, H., et al. (2008). Mouse let-7 miRNA populations exhibit RNA editing that is constrained in the 5’-seed/ cleavage/anchor regions and stabilize predicted mmu-let-7a:mRNA duplexes. Genome Res 18, 1571–1581.

Reimand, J., Isserlin, R., Voisin, V., Kucera, M., Tannus-Lopes, C., Rostamianfar, A., Wadi, L., Meyer, M., Wong, J., Xu, C.J., et al. (2019). Pathway enrichment analysis and visualization of omics data using g:Profiler, GSEA, Cytoscape and EnrichmentMap. Nature protocols 14, 482–517.

Richards, S., Aziz, N., Bale, S., Bick, D., Das, S., Gastier-Foster, J., Grody, W.W., Hegde, M., Lyon, E., Spector, E., et al. (2015). Standards and guidelines for the interpretation of sequence variants: a joint consensus recommendation of the American College of Medical Genetics and Genomics and the Association for Molecular Pathology. Genetics in Medicine 17, 405–424.

Saitsu, H., Kato, M., Mizuguchi, T., Hamada, K., Osaka, H., Tohyama, J., Uruno, K., Kumada, S., Nishiyama, K., Nishimura, A., et al. (2008). De novo mutations in the gene encoding STXBP1 (MUNC18-1) cause early infantile epileptic encephalopathy. Nat Genet 40, 782–788.

Sanders, S.J., Murtha, M.T., Gupta, A.R., Murdoch, J.D., Raubeson, M.J., Willsey, A.J., Ercan-Sencicek, A.G., DiLullo, N.M., Parikshak, N.N., Stein, J.L., et al. (2012). De novo mutations revealed by whole-exome sequencing are strongly associated with autism. Nature 485, 237–U124.

Sanders, S.J., Xin, H., Willsey, A.J., Ercan-Sencicek, A.G., Samocha, K.E., Cicek, A.E., Murtha, M.T., Bal, V.H., Bishop, S.L., Shan, D., et al. (2015). Insights into Autism Spectrum Disorder Genomic Architecture and Biology from 71 Risk Loci. Neuron 87, 1215–1233.

Satterstrom, F.K., Kosmicki, J.A., Wang, J., Breen, M.S., De Rubeis, S., An, J.-Y., Peng, M., Collins, R., Grove, J., Klei, L., et al. (2019). Large-scale exome sequencing study implicates both developmental and functional changes in the neurobiology of autism. bioRxiv, 484113.

Seto, T., Hamazaki, T., Nishigaki, S., Kudo, S., Shintaku, H., Ondo, Y., Shimojima, K., and Yamamoto, T. (2017). A novel CASK mutation identified in siblings exhibiting developmental disorders with/without microcephaly. Intractable Rare Dis Res 6, 177–182.

Shannon, P., Markiel, A., Ozier, O., Baliga, N.S., Wang, J.T., Ramage, D., Amin, N., Schwikowski, B., and Ideker, T. (2003). Cytoscape: A software environment for integrated models of biomolecular interaction networks. Genome Res 13, 2498–2504.

Shomron, N., Malca, H., Vig, I., and Ast, G. (2002). Reversible inhibition of the second step of splicing suggests a possible role of zinc in the second step of splicing. Nucleic Acids Res 30, 4127–4137.

Srivastava, S., McMillan, R., Willis, J., Clark, H., Chavan, V., Liang, C., Zhang, H., Hulver, M., and Mukherjee, K. (2016). X-linked intellectual disability gene CASK regulates postnatal brain growth in a non-cell autonomous manner. Acta Neuropathol Commun 4, 30.

Stamberger, H., Nikanorova, M., Willemsen, M.H., Accorsi, P., Angriman, M., Baier, H., Benkel-Herrenbrueck, I., Benoit, V., Budetta, M., Caliebe, A., et al. (2016). STXBP1 encephalopathy A neurodevelopmental disorder including epilepsy. Neurology 86, 954–962.

Stamouli, S., Anderlid, B.M., Willfors, C., Thiruvahindrapuram, B., Wei, J., Berggren, S., Nordgren, A., Scherer, S.W., Lichtenstein, P., Tammimies, K., et al. (2018). Copy Number Variation Analysis of 100 Twin Pairs Enriched for Neurodevelopmental Disorders. Twin research and human genetics : the official journal of the International Society for Twin Studies 21, 1–11.

Stevenson, D., Laverty, H.G., Wenwieser, S., Douglas, M., and Wilson, J.B. (2000). Mapping and expression analysis of the human CASK gene. Mammalian genome : official journal of the International Mammalian Genome Society 11, 934–937.

Stritt, S., Nurden, P., Turro, E., Greene, D., Jansen, S.B., Westbury, S.K., Petersen, R., Astle, W.J., Marlin, S., Bariana, T.K., et al. (2016). A gain-of-function variant in DIAPH1 causes dominant macrothrombocytopenia and hearing loss. Blood 127, 2903–2914.

Subramanian, A., Tamayo, P., Mootha, V.K., Mukherjee, S., Ebert, B.L., Gillette, M.A., Paulovich, A., Pomeroy, S.L., Golub, T.R., Lander, E.S., et al. (2005). Gene set enrichment analysis: a knowledge-based approach for interpreting genome-wide expression profiles. Proc Natl Acad Sci U S A 102, 15545–15550.

Sudhof, T.C. (2012). The presynaptic active zone. Neuron 75, 11–25.

Tammimies, K., Marshall, C.R., Walker, S., Kaur, G., Thiruvahindrapuram, B., Lionel, A.C., Yuen, R.K., Uddin, M., Roberts, W., Weksberg, R., et al. (2015). Molecular Diagnostic Yield of Chromosomal Microarray Analysis and Whole-Exome Sequencing in Children With Autism Spectrum Disorder. Jama 314, 895–903.

Tukiainen, T., Villani, A.C., Yen, A., Rivas, M.A., Marshall, J.L., Satija, R., Aguirre, M., Gauthier, L., Fleharty, M., Kirby, A., et al. (2017). Landscape of X chromosome inactivation across human tissues. Nature 550, 244–248.

Uddin, M., Woodbury-Smith, M., Chan, A., Brunga, L., Lamoureux, S., Pellecchia, G., Yuen, R.K.C., Faheem, M., Stavropoulos, D.J., Drake, J., et al. (2017). Germline and somatic mutations in STXBP1 with diverse neurodevelopmental phenotypes. Neurol Genet 3, e199.

Uhlin, E., Rönnholm, H., Day, K., Kele, M., Tammimies, K., Bölte, S., and Falk, A. (2017). Derivation of human iPS cell lines from monozygotic twins in defined and xeno free conditions. Stem cell research 18, 22–25.

Wang, G.S., Hong, C.J., Yen, T.Y., Huang, H.Y., Ou, Y., Huang, T.N., Jung, W.G., Kuo, T.Y., Sheng, M., Wang, T.F., et al. (2004). Transcriptional modification by a CASK-interacting nucleosome assembly protein. Neuron 42, 113–128.

Wincent, J., Kolbjer, S., Martin, D., Luthman, A., Amark, P., Dahlin, M., and Anderlid, B.M. (2015). Copy Number Variations in Children with Brain Malformations and Refractory Epilepsy. American Journal of Medical Genetics Part A 167, 512–523.

Yuen, R.K.C., Merico, D., Bookman, M., Howe, J.L., Thiruvahindrapuram, B., Patel, R.V., Whitney, J., Deflaux, N., Bingham, J., Wang, Z.Z., et al. (2017). Whole genome sequencing resource identifies 18 new candidate genes for autism spectrum disorder. Nat Neurosci 20, 602-+.

Zweier, C., de Jong, E.K., Zweier, M., Orrico, A., Ousager, L.B., Collins, A.L., Bijlsma, E.K., Oortveld, M.A., Ekici, A.B., Reis, A., et al. (2009). CNTNAP2 and NRXN1 are mutated in autosomal-recessive Pitt-Hopkins-like mental retardation and determine the level of a common synaptic protein in Drosophila. Am J Hum Genet 85, 655–666.

